# PKCβ facilitates leukemogenesis in chronic lymphocytic leukaemia by promoting constitutive BCR-mediated signaling

**DOI:** 10.1101/2022.09.04.506520

**Authors:** Jodie Hay, Anuradha Tarafdar, Ailsa Holroyd, Hothri A. Moka, Karen M. Dunn, Alzahra Alshayeb, Bryony H. Lloyd, Jennifer Cassels, Natasha Malik, Ashfia F. Khan, IengFong Sou, Jamie Lees, Hassan N. B. Almuhanna, Nagesh Kalakonda, Joseph R. Slupsky, Alison M. Michie

## Abstract

B cell antigen receptor (BCR) signaling competence is critical for pathogenesis of chronic lymphocytic leukemia (CLL). Defining key proteins that facilitate these networks aid in the identification of targets for therapeutic exploitation. We previously demonstrated that reduced PKCα function in mouse hematopoietic stem/progenitor cells (HPSCs) resulted in PKCβII upregulation and generation of a poor-prognostic CLL-like disease. Here, *prkcb* knockdown in HSPCs leads to reduced survival of PKCα-KR-expressing CLL-like cells, concurrent with reduced expression of the leukemic markers CD5 and CD23. SP1 promotes elevated expression of *prkcb* in PKCα-KR expressing cells enabling leukemogenesis. Global gene analysis revealed an upregulation of genes associated with B cell activation in PKCα-KR expressing cells, coincident with upregulation of PKCβII: supported by activation of key signaling hubs proximal to the BCR and elevated proliferation. Ibrutinib (BTK inhibitor) or enzastaurin (PKCβII inhibitor) treatment of PKCα-KR expressing cells and primary CLL cells showed similar patterns of Akt/mTOR pathway inhibition, supporting the role for PKCβII in maintaining proliferative signals in our CLL mouse model. Ibrutinib or enzastaurin treatment also reduced PKCα-KR-CLL cell migration towards CXCL12. Overall, we demonstrate that PKCβ expression facilitates leukemogenesis and identify that BCR-mediated signaling is a key driver of CLL development in the PKCα-KR model.

**Statement of Significance:** PKCβ facilitates leukemogenesis of CLL, driven through an SP1-regulated transcriptional program and promotes BCR signaling. Thus far, PKCβ is the only kinase within the BCR signaling pathway, a key pathway in driving CLL pathogenesis, implicated in the generation of neoplastic B lineage cells.

## Introduction

Intracellular signals transmitted through the B cell antigen receptor (BCR) regulate B cell differentiation, survival and proliferation at distinct stages of development, maturation and activation, and play a critical role in driving the pathogenesis of chronic lymphocytic leukemia (CLL) [1]. The BCR is a critical prognostic marker of CLL, with sequences aligning closely with germline (>98% similarity; unmutated IgHV) generally associated with poor prognostic CLL cases [2]. The structure of the BCR appears to be related to its ability to transmit and activate intracellular signaling networks more efficiently, thus promoting CLL cell survival and proliferation [3]. BCR crosslinking promotes membrane localization and activation of a number of cytoplasmic protein kinases including Lyn and spleen tyrosine kinase (SYK), and activates Bruton’s tyrosine kinase (BTK), triggering the activation of multiple key signaling pathways and transcription factors that regulate B cell proliferation, differentiation and apoptosis in a cell maturation-dependent manner [4].

Membrane-recruited BTK undergoes auto-phosphorylation, and then activates phospholipase C (PLCγ2) [5] which further catalyzes the hydrolysis of phosphatidylinositol 4,5-bisphosphate (PIP_2_) into diacylglycerol (DAG) and inositol trisphosphate (IP_3_). Intracellular calcium levels increase as a result of IP_3_ production, thus providing the cofactors required for classical protein kinase C (PKC: PKCα, βI/II and γ) isoform activation [6]. PKC isoforms, specifically PKCβII in B cells, activate survival pathways through NF-κB-mediated signaling [7], thus linking proximal BCR-mediated signals with downstream pathways. CLL cells exhibit a deregulated PKC isoform expression profile: upregulation of PKCβII, PKCε, PKCζ and downregulation of PKCα and PKCβI compared with normal B cells, with the increased levels of PKCβII expression correlating with poor prognostic outcome for CLL patients [8]. Furthermore, ZAP-70 recruits PKCβII to lipid rafts in CLL cells, which activates and enhances the translocation of PKCβII to the mitochondria where it phosphorylates the anti-apoptotic protein BCL2 [9].

PKCβ has been shown to play an essential role in the regulation of leukemia development in the TCL1 mouse model of CLL, as indicated by the abrogation of leukemic cells in PKCβ KO mice [10]. We developed a CLL-like mouse model, by introducing a kinase inactive PKCα (PKCα-KR) construct in mouse hematopoietic stem/progenitor cells (HSPCs), which resulted in the development of an aggressive subset of CLL *in vitro* and *in vivo*, exhibiting an upregulation of ZAP-70, enhanced proliferation and increased tumor load in the lymphoid organs [11]. Analysis of the PKC isoform profile in the PKCα-KR-transformed cells revealed that, similar to that seen in primary CLL samples, PKCβII expression was upregulated, and this occurred at later stages of disease development [12]. Here we determine the importance of PKCβ expression in disease development in our mouse model, analyze the mechanisms that promote PKCβ upregulation, and delineate the signaling pathways that occur downstream of BTK/PKCβ, which may have particular importance in developing future therapies for B cell malignancies.

## Materials and Methods

### Primary Cells and Cell lines

HSPCs obtained from fetal liver (FL) of wild type ICR mice on day 14 of gestation were prepared as described previously, and retrovirally-transduced with a plasmid encoding either empty vector control (MIEV) or MIEV-PKCα-KR (PKCα-KR) with bicistronic expression of GFP. Maintenance of the MIEV constructs during the course of the cultures was followed by assessing GFP expression using flow cytometry [11]. Immediately prior to analysis, MIEV and PKCα-KR cells were removed from OP9 monolayers and placed on empty 6 well plates for 2 hr to remove carry-over OP9 cells. All cell lines were routinely tested for mycoplasma contamination. All mice were maintained at the University of Glasgow Central Research Facilities under standard animal housing conditions in accordance with local and home office regulations (PD6C67A47). Primary CLL cells, obtained from patients that had given informed consent, were isolated as described previously [13] and cryopreserved for future use. The studies were approved by the West of Scotland Research Ethics Service, NHS Greater Glasgow and Clyde (UK) and all work was carried out in accordance with the approved guidelines (REC Ref: 20/WS/0066). Clinical details of patients used in these studies are presented in Supplementary Table 1; none of the patients had received chemotherapy within the preceding 3 months.

### Surface and Intracellular staining and flow cytometric analysis

Flow cytometry reagents used are detailed in Supplementary Table 2. After drug treatment/stimulation as indicated, MIEV and PKCα-KR cells (≥1 × 10^6^ cell/condition) were stained as described previously [11]. Cells were acquired using a FACSCantoII flow cytometer with BD FACSDiva software and analyzed using FlowJo (Tree Star Inc., Ashland, USA) software.

### Cell Stimulation/drug treatment

MIEV and PKCα-KR cells (≥1×10^6^/condition) were treated with 1 μM ibrutinib (IB) or 20 μM enzastaurin (enza) or vehicle control in 10% FBS/αMEM media as previously published [12]. Primary CLL cells were pre-incubated with 1 μM IB, 10 μM Enza or vehicle control in 10% FBS/RPMI media on ice and then stimulated with 10 μg/ml F(ab’)_2_ fragment anti-human IgM (Stratech Scientific Ltd.) to crosslink the BCR (BCR-XL) at 37°C and then analyzed.

### Cell counting beads

To compare cell counts between different co-culture conditions, a set volume of CountBright beads (BDBiosciences) was added to each PKCα-KR sample as indicated, prior to flow cytometry. After FSC/SSC gating on the beads, 2000 beads were acquired together with a variable cell number to enable a relative cell number to be calculated between samples [14].

### Proliferation assays

MIEV and PKCα-KR cells (≥1 × 10^6^/condition) were labelled with CellTrace Violet (CTV; 2 μM) using CellTrace™ Cell Proliferation Kits (Life Technologies) as described previously [14]. Cells were analyzed on the FACSCantoII flow cytometer. Results are expressed relative to the mean fluorescence intensity (MFI) of CTV in no drug control (NDC) cells at 48hr.

### Lentiviral knockdown of prkcb

Lentiviral knockdown vectors targeting mouse *prkcb* and non-targeting control (SCR) were purchased from Sigma-Aldrich (St. Louis, MO) in the pLKO.1 backbone. Lentiviral particles were generated using a three-plasmid HEK293T lentiviral protocol, as described previously [15]. Briefly, HEK293 cells were transfected overnight using CaCl_2_ and 2xHBS containing pHIV1 and VSV-G as accessory plasmids. Virus was collected in DMEM supplemented with 20% FCS and freshly-isolated HSPCs (d0) were cultured in viral medium supplemented with 4 μg/ml polybrene and IL7/Flt3 (10 ng/ml each). This was repeated 4 times over 48 hr. After transduction the cells were washed and re-suspended in complete media. Cell sorting for GFP^+^CD19^+^CD45^+^CD11b^-^ cells was performed using FACSAria™ III cell sorter (BD Biosciences) on day 7; MIEV and PKCα-KR were then introduced by retroviral transduction on day 8 as described previously [11]. *qRT-PCR:* RNA was extracted from MIEV-or PKCα-KR-transduced cells from day 17-23 co-cultures using RNeasy mini kit (Qiagen, UK) according to manufacturer’s protocol. Synthesis of cDNA was performed using SuperScript ® III Reverse transcriptase (Life Technologies) as per manufacturer’s protocol. Real-time PCR (RT-PCR) was conducted using the TaqMan PCR Master Mix (Applied Biosystems), with glyceraldehyde-3-phosphate dehydrogenase (*gapdh*) used as a reference control. Inventoried primers and probes and PCR buffers were purchased from Applied Biosystems unless otherwise stated. *Egr1* and *Btk* primers were designed and optimized, using TATA-box binding protein (*tbp*) as a reference control (Supplementary Table 3). For each PCR reaction 1 μl cDNA was used followed by appropriate primer master mix containing 0.25 μM forward, 0.25 μM reverse primers and 2xSyber Green PCR Mastermix (ThermoFisher Scientific). Technical triplicates of each PCR reaction were performed. qRT PCR was performed on the 7900HT Fast Real Time PCR System (Applied Biosystems): The cycle condition for 40 cycles is as follows: 95°C 2 min, 40 cycles of 95°C 5 s, 60°C10 s, 72°C 5-20 s. NDC or MIEV was used as the calibrator and data was analyzed using the 2^-ΔΔCT^ relative quantification method.

### Chromatin immunoprecipitation (ChIP)

MIEV- and PKCα-KR-transduced cells were co-cultured on OP9 cells until day 20-23, as described above. The cells were harvested and IP was performed on the sonicated chromatin material using a ChIP grade antibody -anti-SP1 antibody (Cell Signaling Technologies, Clone D4C3) or IgG (negative control), as described previously [16]. The primer sets were designed on regions flanking three SP1 binding regions within the *prkcb* promoter (Figure 2A; Supplementary Table 4).

### Microarrays

MIEV-or PKCα-KR-transduced HSPCs were co-cultured on OP9 cells in the presence of IL-7 for 17-23 days (late co-culture) and total RNA was isolated using an RNeasy kit (Qiagen, Manchester, UK) from five independent co-cultures. The RNA was quantified using a Nanodrop ND-1000 Spectrophotometer (ThermoScientific, Waltham, MA). Screening of the RNA samples was performed against Affymetrix™ GeneChip® Mouse Gene 1.0 ST Array. Background correction and normalization of generated CEL files were done using Robust Multichip Average (RMA) in Partek Genomics Suite 7.19.1125 (St Louis, MI). Differentially expressed genes, PKCα-KR vs. MIEV, were detected by one-way analysis of variance. Significant genes (n= 2836), were identified with a fold change ±1.2 and p-value <0.05, with 1513 up- and 1323 downregulated in PKCα-KR vs. MIEV. Microarray data are available at the Gene Expression Omnibus (GEO) repository (GSE185075).

### Western Blots

Cell lysates were prepared in lysis buffer (1% Triton, 1 mM DTT, 2 mM EDTA, 20 mM Tris pH 7.5 containing complete protease inhibitor, and PhosStop; Roche) from early (day 6-10) or late (day 17-23) MIEV and PKCα-KR cells, or primary CLL cells, with or without drug treatment as indicated and quantified using the Bradford Assay (ThermoFisher Scientific). The antibodies used are described in Supplementary Table 5, and Western blotting was performed as described previously [14]. Images were developed using SRX 101A film processor.

### Migration assay

MIEV-or PKCα-KR-transduced cells (2 × 10^5^) were incubated in 100 μl RPMI-1640/0.5% BSA media in the presence or absence of drugs (as indicated) for 30 min prior to the assay. Cells were then transferred to the upper chamber of a 6.5-mm diameter Transwell culture insert (Costar^®^, Fisher Scientific UK.) and placed into wells containing 600 μl media supplemented with 150 ng/ml CXCL12. The cells were incubated for 4 hr at 37°C. Thereafter, three 150 μl aliquots were removed from each lower chamber for counting by flow cytometry. The total number of events acquired during 20 s on high flow setting was recorded for each aliquot.

### Statistics

p values were determined by students paired or unpaired *t*-test or mixed model ANOVA on a minimum of at least 3 biological replicates to compare data using GraphPad Prism 6 software (GraphPad Software Inc.) as indicated; *, **, *** and **** represent *p* <0.05, 0.01, 0.001 and < 0.0001 respectively. Biological replicates were derived from individual cell culture conditions from distinct biological samples. Sample sizes were chosen based on the effect size observed within initial experiments, and based on the heterogeneity of the biological samples. Results are presented as mean ± standard error of mean of biological replicates (SEM), with the number of replicates performed highlighted in the figure legends.

## Results

### PKCβ promotes leukemogenesis in the PKCα-KR CLL-like mouse model

We and others have previously demonstrated that PKCβII is upregulated in CLL mouse models (Eμ-Tcl-1 and PKCα-KR; [12]) and in primary CLL samples, particularly in poor prognostic samples suggesting a role in CLL pathogenesis [8]. To extend this research we addressed the importance of *prkcb* expression during the initiation of the CLL-like disease in our PKCα-KR mouse model. Performing *prkcb* shRNA knockdown (KD) experiments prior to retroviral transduction with PKCα-KR revealed that generation of the leukemic phenotype was negatively impacted upon reduction of *prkcb* expression. There was a significant reduction in surface expression of the leukemic markers characteristic of this CLL model, CD23 and CD5 (Figure 1A&B), as well as CD45 expression. An additional feature of PKCα-KR-transduced cells is an increased cellular count in the cultures, which was significantly reduced, together with elevated apoptosis in *prkcb* KD conditions (Figure 1C&D). A significant reduction in PKCβ mRNA levels was confirmed in cultured cells transduced with sh-*prkcb*, compared with SCR controls (Figure 1E). While the reduced cell numbers generated with PKCα-KR-transduced *prkcb* KD cells precluded us from being able to test whether CLL-like development was blocked *in vivo*, these results indicate that PKCβ plays an important role in the initiation of CLL in our mouse model, supporting previous findings in the Eμ-Tcl-1 CLL mouse model [10].

**Figure 1.**
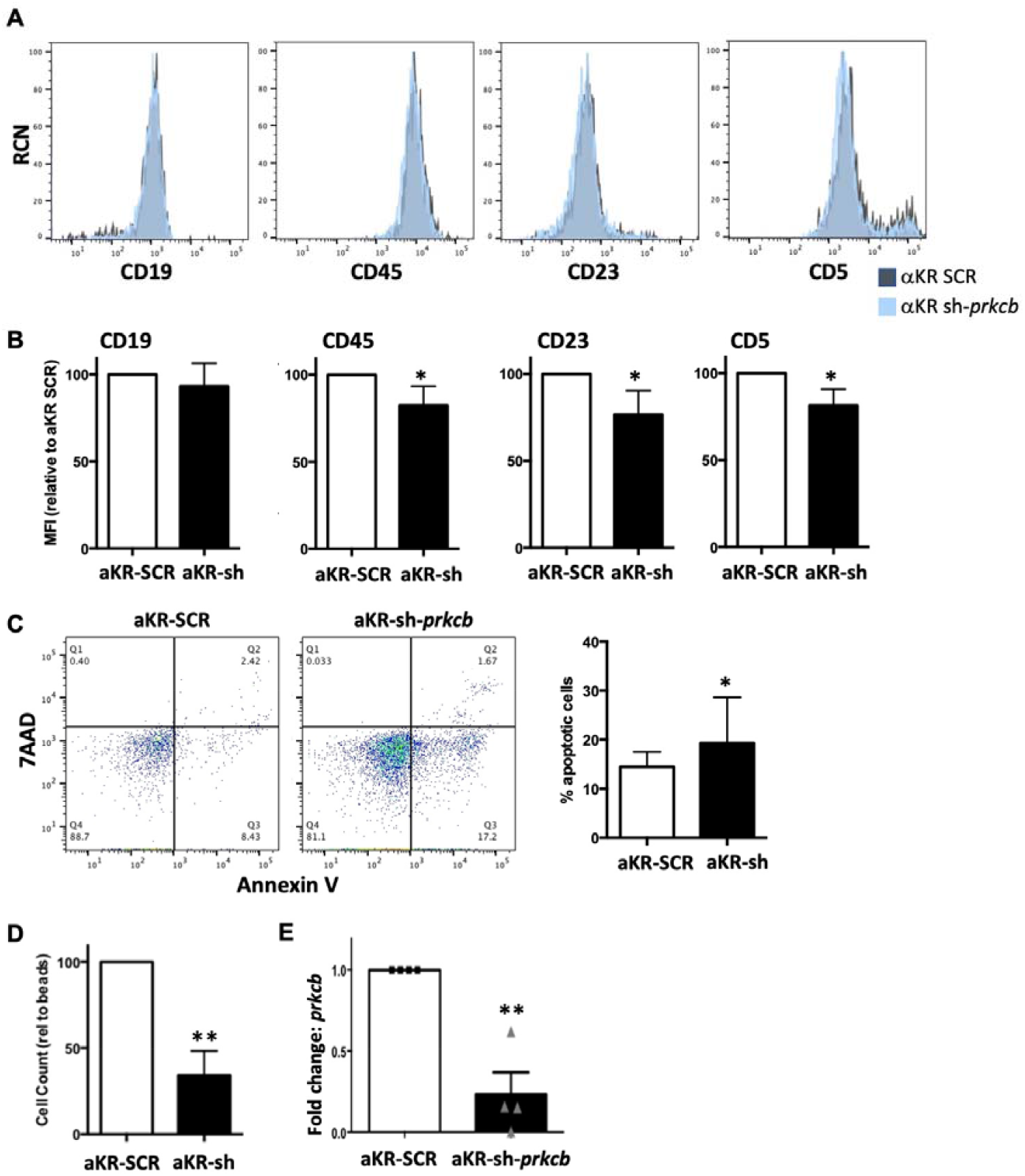
Reduction in PKCβ expression inhibits the initiation of PKCα-KR-mediated CLL development. Knockdown of *prkcb* (or scrambled (SCR) control) was performed in HSPC cells with 24 hr of isolation from the mouse, prior to PKCα-KR (αKR) retroviral transduction at d7. Resultant cells were co-cultured with OP9 in the presence of cytokines for up to 35 days. Phenotypic characterization of the cells was carried out by flow cytometry analyzing: A. CD19, CD45, CD23 and CD5. Representative histogram plots are shown, gated on FSC/SSC and GFP+ cells comparing αKR sh-*prkcb* cells with αKR SCR cells; B. Average MFIs of CD19, CD45, CD23 and CD5 surface markers are shown relative to αKR-SCR cultures (n=5); C. Apoptosis was determined by AnnV/7AAD staining. A representative dot plot shows Annexin V vs. 7AAD staining in αKR SCR cells and αKR sh-*prkcb*. The graph shows the percentage apoptotic (AnnV+) cells present in cultures post d14 (n=5); D. Cell counts were performed of each co-culture using flow cytometry using counting beads, shown relative to a set bead number acquired (n=5 biological replicates); E. qPCR analysis of *prkcb* expression. *Gapdh* was used as the reference gene and SCR-PKCα-KR-transduced cells were used as a calibrator (n=4 biological replicates). All experiments shown are representative of n≥3 biological replicates. Paired student t test with Wilcoxon matched pairs signed rank test was used to analyze the data.

### Sp1 regulates similar transcriptional networks in the PKCα-KR CLL-like cells and primary human CLL samples

The transcription factor SP1 plays a central role in *PRKCB* gene regulation in human CLL [17]. We were interested in determining whether the *prkcb* gene was similarly regulated by Sp1 in our CLL mouse model. The *prkcb* promoter contains three Sp1 binding regions (Figure 2A). ChIP analysis revealed that Sp1 binding to the *prkcb* promoter was significantly upregulated in PKCα-KR transduced CLL-like cells at all three Sp1 binding sites, compared with MIEV cells (Figure 2B). This data was supported with experiments using mithramycin (mtm), an anti-cancer antibiotic that selectively binds G-C rich DNA sequences to inhibit RNA and DNA polymerases and globally displace Sp1 [18]. Treatment of PKCα-KR cells with 200 nM mtm significantly reduced Sp1 binding activity at all regions assessed on the *prkcb* promoter (Figure 2C). Mtm treatment also resulted in a downregulation of PKCβII protein expression specifically in PKCα-KR cells, which was elevated in untreated cells compared to MIEV cells, although the mtm-mediated downregulation of PKCβII expression did not reach significance (Figure 2D, Supplementary Figure 1). Assessing the wider impact of mtm treatment on gene transcription in our PKCα-KR cells revealed a reduction in expression not only of *prkcb*, but also *Sp1* itself and *Bcl2, Vegfa, Blnk* and *Lef1* upon treatment with 200 nM mtm, compared to no drug control (NDC) cells (Figure 2E), a result similar to that seen in human CLL samples (Supplementary Table 6). Taken together, these data suggest a similar regulation of Sp1-mediated transcription networks between poor prognostic human CLL cells and PKCα-KR CLL cells.

**Figure 2.**
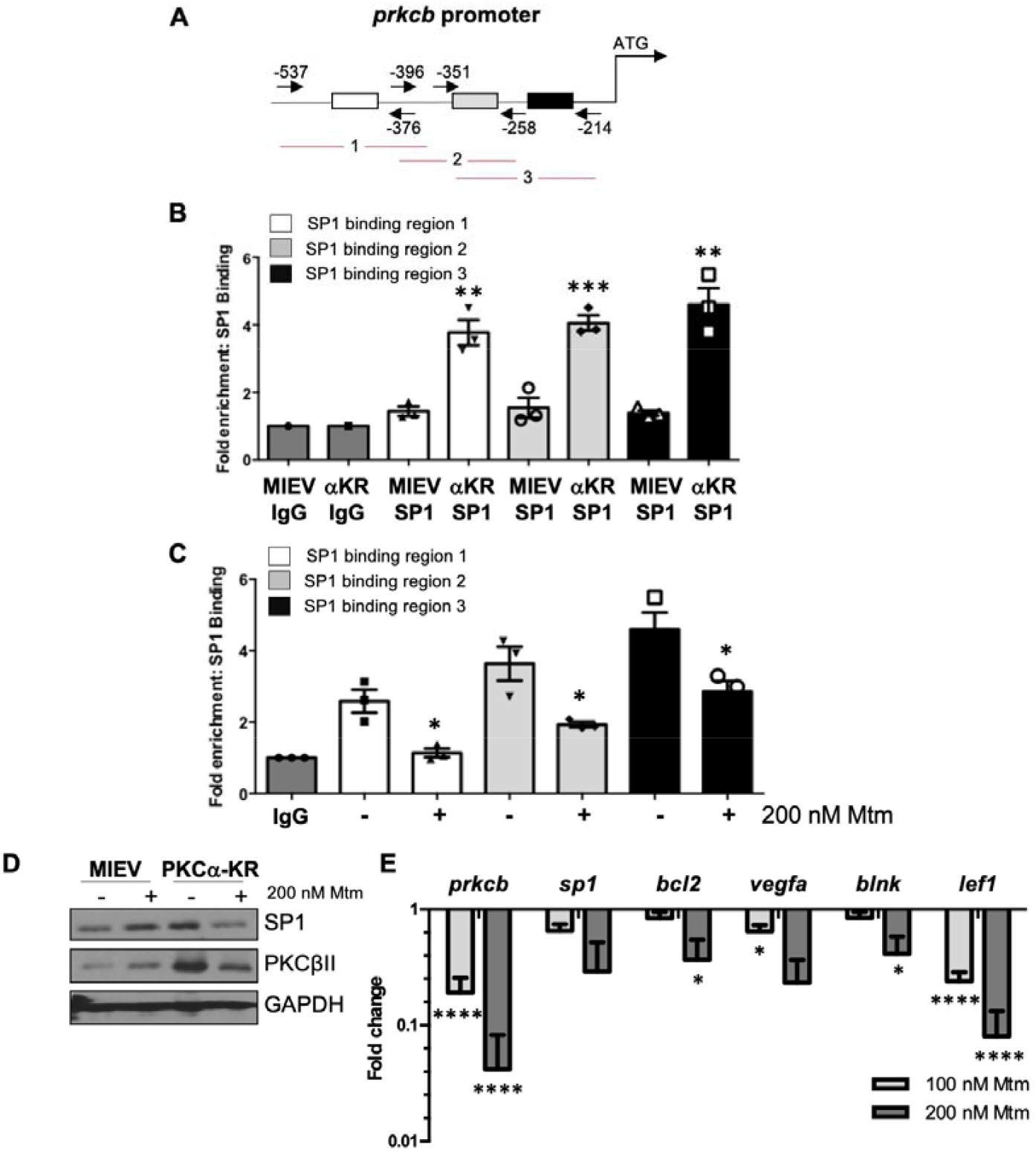
Sp1 binding is upregulated in the *prkcb* promoter of PKCα-KR transduced CLL like cells. A. Diagram showing Sp1 binding sites at the *prkcb* promoter, the primer locations and the three products; B. ChIP analysis was performed in MIEV and PKCα-KR cells at the late stage of co-culture (> d17) to determine Sp1 binding occupancy at the *prkcb* promoter (average of n=3 biological replicates); C. ChIP analysis was performed in PKCα-KR cells treated with 200 nM mithramycin (mtm) for 12 hr to determine Sp1 binding at the *prkcb* promoter (average of n=3 biological replicates); D. The expression levels of Sp1 and PKCβII were determined by Western blotting in MIEV and PKCα-KR cells treated with 200 nM mtm (representative blot shown of n=3 experiments; densitometry of the blots shown in Supplementary Figure 1). E. qPCR analysis of *prkcb, sp1, bcl2, vegfa, blnk, lef1* genes in PKCα-KR cells upon treatment with mtm (average of n=2/3 biological replicates, an average of technical duplicates). *Gapdh* was used as the reference gene and normalized to NDC. Unpaired student t tests (B & C) or one way ANOVA (E) were used to analyze the data (where biological triplicates were performed).

### BCR signaling components are deregulated in PKCα-KR CLL-like cells

To investigate global gene expression profiles in the PKCα-KR CLL mouse model within the timeline of PKCβII upregulation, microarrays were performed comparing MIEV- and PKCα-KR-transduced cells at the late stage of B cell transformation (post d17 of co-culture after retroviral transduction at d0 [12]). Analysis of the dataset revealed differentially regulated gene expression between the MIEV- and PKCα-KR-transduced cells (Supplementary Figure 2), and enrichment of pathways that were significantly deregulated, with immune system processes identified as the most enriched group of pathways (Table 1 & Supplemental Figure 3). This analysis indicated an upregulation of genes allied to B cell activation in PKCα-KR-transduced cells, and given the importance of BCR signaling in the pathology of CLL, we chose to validate this pathway in our CLL mouse model (Figure 3). Deregulation of the BCR signaling in PKCα-KR-transduced cells, summarized in Figure 3A, is highlighted by a number of features including, CD45 and CD19 upregulation in the CLL-like PKCα-KR cells (indicated in pink; previously confirmed by flow cytometry [12]), together with an upregulation of signaling components proximal to BCR-mediated signaling (*Lyn, Btk, Blnk*) while *Syk, Pten* and *Pdk1* are downregulated (indicated in green) compared to MIEV control cells. The expression of transcription factors was deregulated with PKCα-KR cells showing an upregulation of *Bcl6, Egr1, Elk1, NFκB* and *Foxo1*, while *Creb* and *Tcf1* were downregulated compared to MIEV control cells in the microarray datasets. The upregulation of *Egr1* and *Btk* in the late PKCα-KR cultures was confirmed by qPCR (Figure 3B). In support of the data shown in Figure 2E, *Prkcb* and *Lef1* expression patterns aligned with the *Sp1* expression, with downregulation of these genes in early PKCα-KR cultures, and significant upregulation in the late PKCα-KR cultures, compared to the respective MIEV cells (Figure 3C).

**Table 1.**
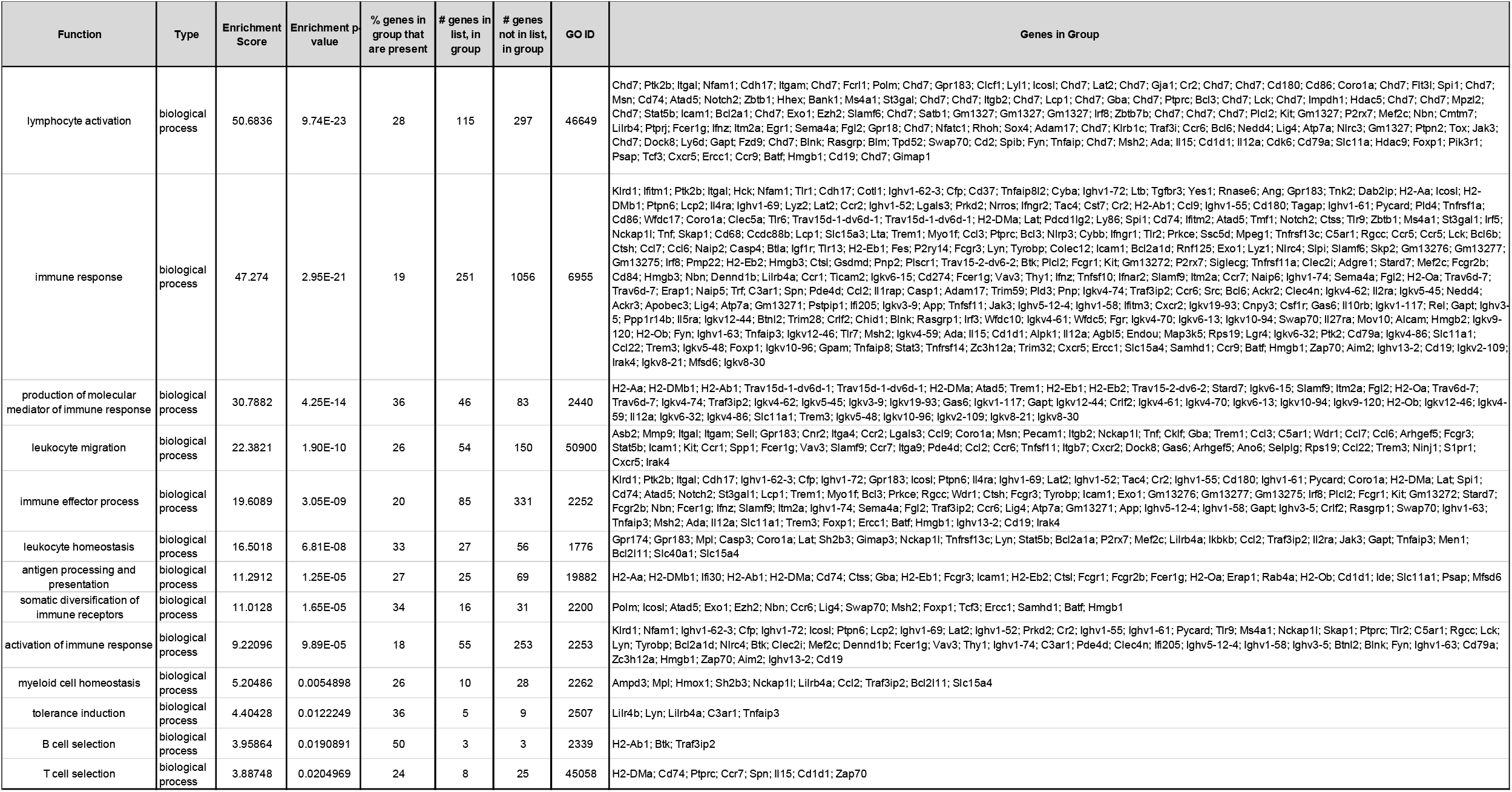
Gene ontology analysis of the significantly altered genes (fold change ±1.2 and p-value <0.05) between late co-culture PKC⍰-KR vs. MIEV cells, identifies the most enriched group of pathways.

**Figure 3.**
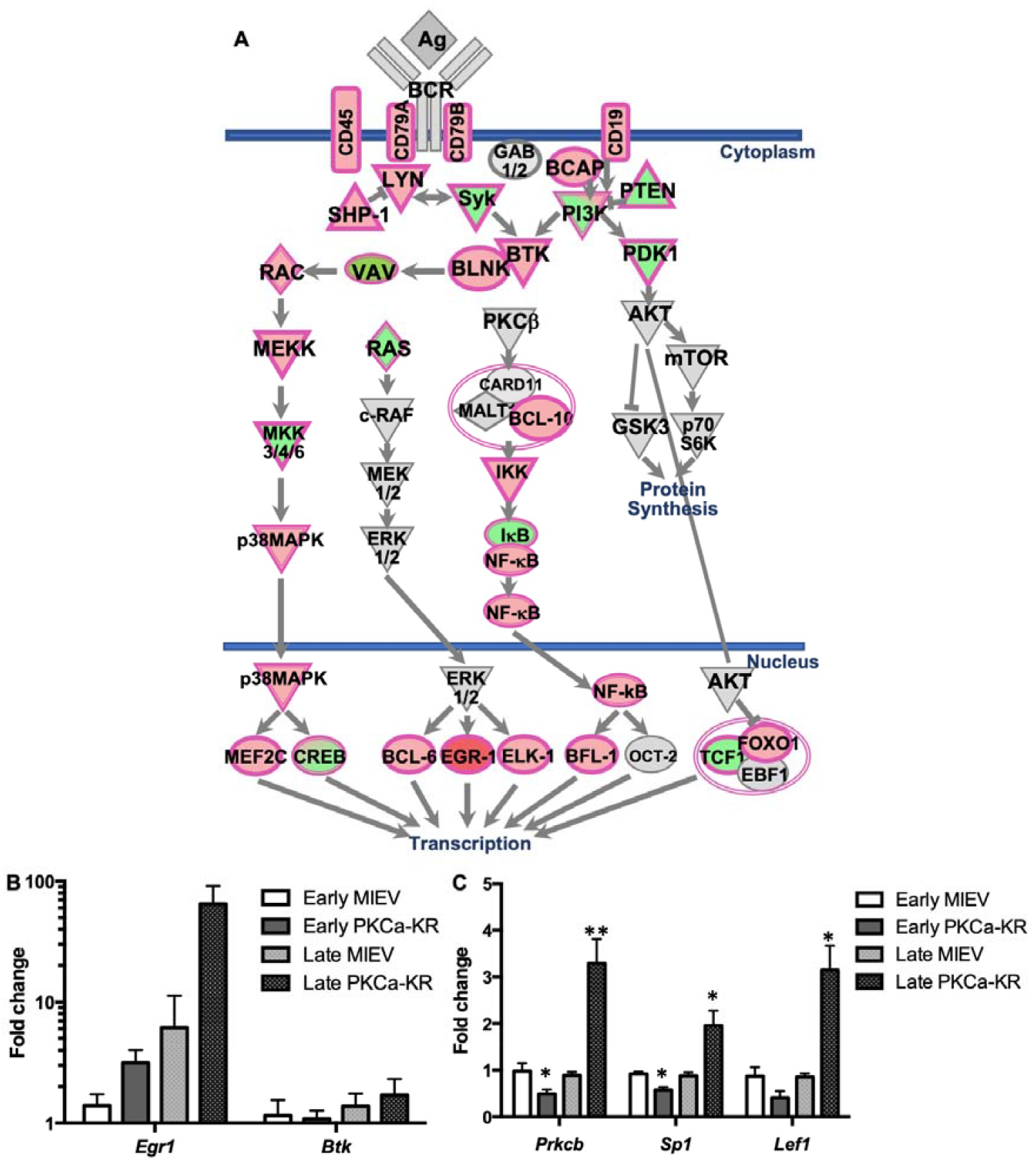
Global gene expression analysis revealed activation of the BCR-mediated signaling pathway in PKCα-KR cells. Global RNA analysis was performed using Affymetrix GeneChip mouse gene 1.0 ST on MIEV and PKCα-KR transduced cells at d17-23 (late) in the B cell transformation co-culture. A. Deregulated components of the BCR pathway in PKCα-KR vs. MIEV cells isolated from late co-cultures (n=5 PKCα-KR, n=5 MIEV) were identified. Significantly up- and down-regulated components are highlighted in pink/red and green respectively; B. qPCR validation of *egr1* and *btk* genes, which were both upregulated in the microarray analysis at the late stage (d15-23) of B cell transformation, compared with the early stage (d6-10). Data are an average of n=2-4 biological replicates, normalized to *tbp*. C. Comparison of *prkcb, sp1* and *lef1* gene expression in the early vs. late stages of B cell transformation co-cultures, determined by qPCR. Data represents n=3-5 biological replicates, normalized to *gapdh*.

Analyzing the expression levels of BCR signaling components demonstrated that key hubs proximal and distal to the BCR were deregulated in the CLL-like PKCα-KR cells compared to MIEV control cells both at the early and late stages of the CLL-like disease. Lyn, c-Myc and Egr1 expression was upregulated, while Lck was downregulated (Figure 4A & C & Supplementary Figure 4 & 5). PKCα expression was significantly downregulated in the PKCα-KR cells, in agreement with our previous work [12]. Furthermore, a significant elevation in pAKT^S473^ and pS6^S235/236^ was observed, indicating an elevation in AKT/mTOR mediated signaling in PKCα-KR cultures (Figure 4A, Supplementary Figure 4). An upregulation in pc-Myc^S62^ was also observed, accompanied by an increase in c-Myc expression in PKCα-KR cells compared with MIEV suggesting a stabilization of c-Myc in PKCα-KR cells, however these modulations did not reach significance (Figure 4C, Supplementary Figure 5). Phospho-flow staining revealed a significant elevation of both BTK^Y551^ and BTK^Y223^ phosphorylation in PKCα-KR compared to MIEV cells (Figure 4B). Taken together these results indicate that PKCα-KR CLL-like cells exhibit upregulated BCR signaling activity.

**Figure 4.**
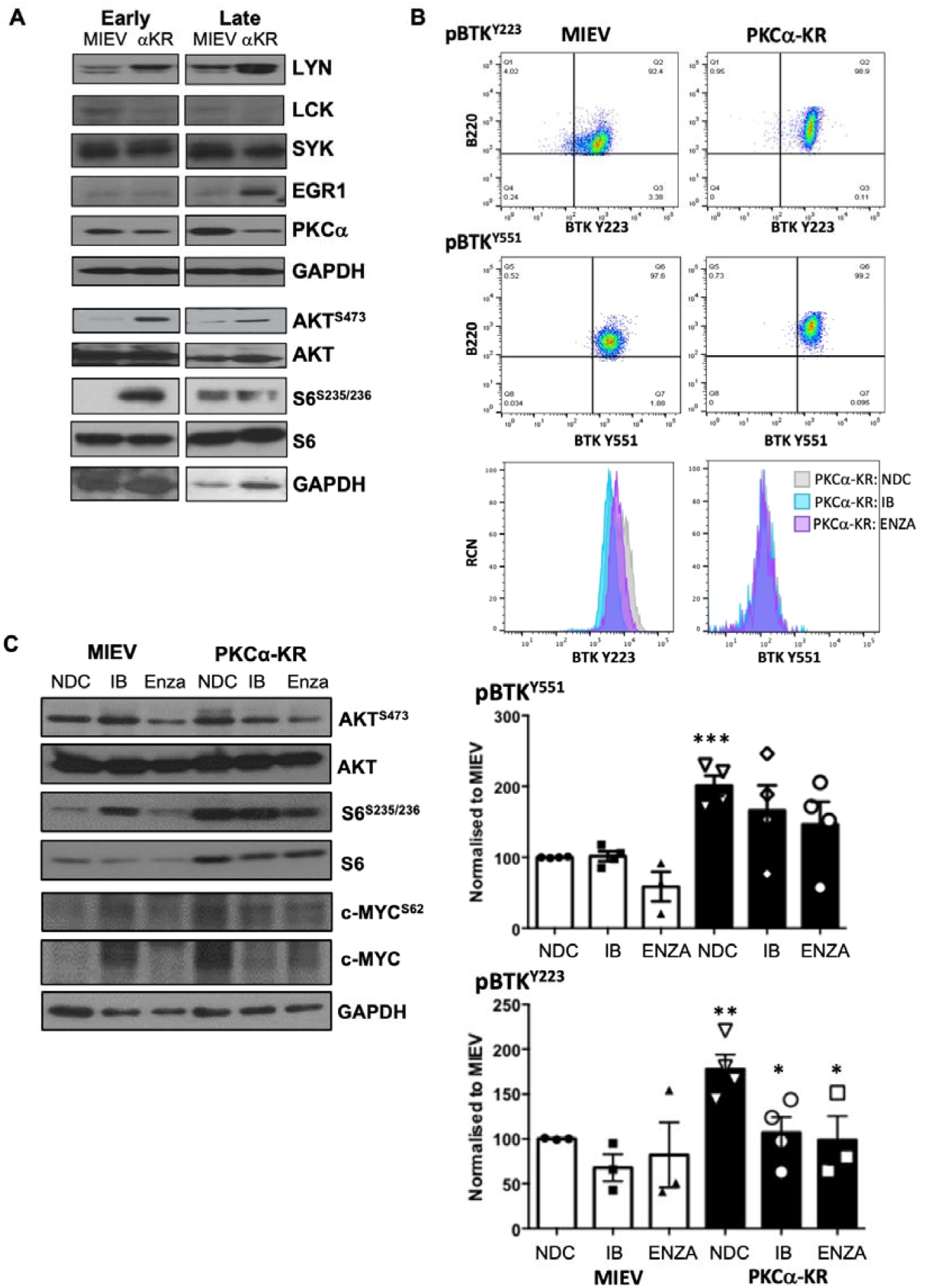
Inhibition of PKCβ activity downregulates key proteins in the BCR signalosome in mouse CLL model. A. The expression/activation status of key hubs proximal and distal to the BCR pathway were analyzed by Western blotting in early and late co-cultures of MIEV and PKCα-KR cells. Representative Western blots are shown. Densitometry of the Western blots for these proteins (n≥4) is shown in Supplementary Figure 4. B. Phospho-flow was used to analyze the levels of BTK^Y551^ and BTK^Y223^ phosphorylation in MIEV (left) and PKCα-KR (right) cells in the presence and absence of either Ibrutinib (IB; 1 μM) or enzastaurin (Enza; 20 μM). Upper panel shows the flow cytometry dot plots of the mouse B lineage cells (B220^+^) vs phospho-BTK, gated on FSC/SSC, while lower histogram plots show the effect of drug treatments on BTK^Y551^ and BTK^Y223^ phosphorylation as overlay (NDC – pale grey; IB – turquoise; ENZA – pink). Lower graphs show the average MFI of phospho-BTK normalized to MIEV no drug control (NDC; n=4 independent experiments; MIEV – white bars, PKCα-KR – black bars). C. MIEV and PKCα-KR cells were treated with either IB or Enza, or left NDC. Representative Western blots are shown identifying the effect on key proteins within the BCR signalosome as indicated. Densitometry of the Western blots for these proteins (n=3) is shown in Supplementary Figure 5.

### PKCα-KR CLL-like and primary human CLL cells share similar responses to BCR-targeted inhibitors

To determine how PKCβ impacted on BCR signaling, MIEV- and PKCα-KR-expressing cells were treated with either BTK inhibitor ibrutinib (IB) or PKCβII inhibitor enzastaurin (Enza). While the drug treatments did not affect the BTK phosphorylation on MIEV B-cells, IB- and Enza-treatment significantly reduced BTK^Y223^ phosphorylation suggesting an inhibition in BTK activity in the PKCα-KR-expressing B cells (Figure 4B). The elevation in S6^S235/236^ phosphorylation in PKCα-KR-expressing cells was inhibited by Enza-treatment. The elevation in c-Myc^S62^ phosphorylation and c-Myc expression in PKCα-KR cells was not significantly affected with drug treatments (Figure 4C, Supplementary Figure 5). These findings suggest that PKCβ plays a vital role in driving the activation of key proteins within the BCR-mediated BTK and AKT/mTOR signaling in PKCα-KR cells.

To establish whether our findings in the PKCα-KR CLL-like model were applicable to primary human CLL samples, we treated human CLL cells with IB or Enza in the presence of F(ab’)_2_ stimulation (BCR-XL) and assessed the phosphorylation/activation status of BTK. Both BTK^Y551^ and BTK^Y223^ phosphorylation levels were significantly downregulated upon PKCβ inhibition with Enza in human CLL cells compared to NDC, while IB treatment led to a significant downregulation of BTK^Y223^ phosphorylation only (Figure 5A). Western blotting of primary CLL cell lysates stimulated with BCR-XL showed an elevation in phosphorylation of AKT^S473^ and pS6^S3235/236^ and an increase in c-Myc expression similar to that noted in PKCα-KR cells. Furthermore, a reduction in AKT^S473^ and pS6^S3235/236^ phosphorylation was more pronounced with enza treatment in the presence of BCR-XL, although this did not reach significance. Enza significantly decreased overall c-Myc^S62^ phosphorylation upon BCR-XL, however coupled with the significant reduction in total c-Myc protein with IB or Enza treatment, this led to a significant increase in pc-Myc^S62^ phosphorylation on the remaining c-Myc in human CLL cells (Figure 5B, Supplementary Figure 6), findings that mirrored trends in PKCα-KR CLL-like cells.

**Figure 5.**
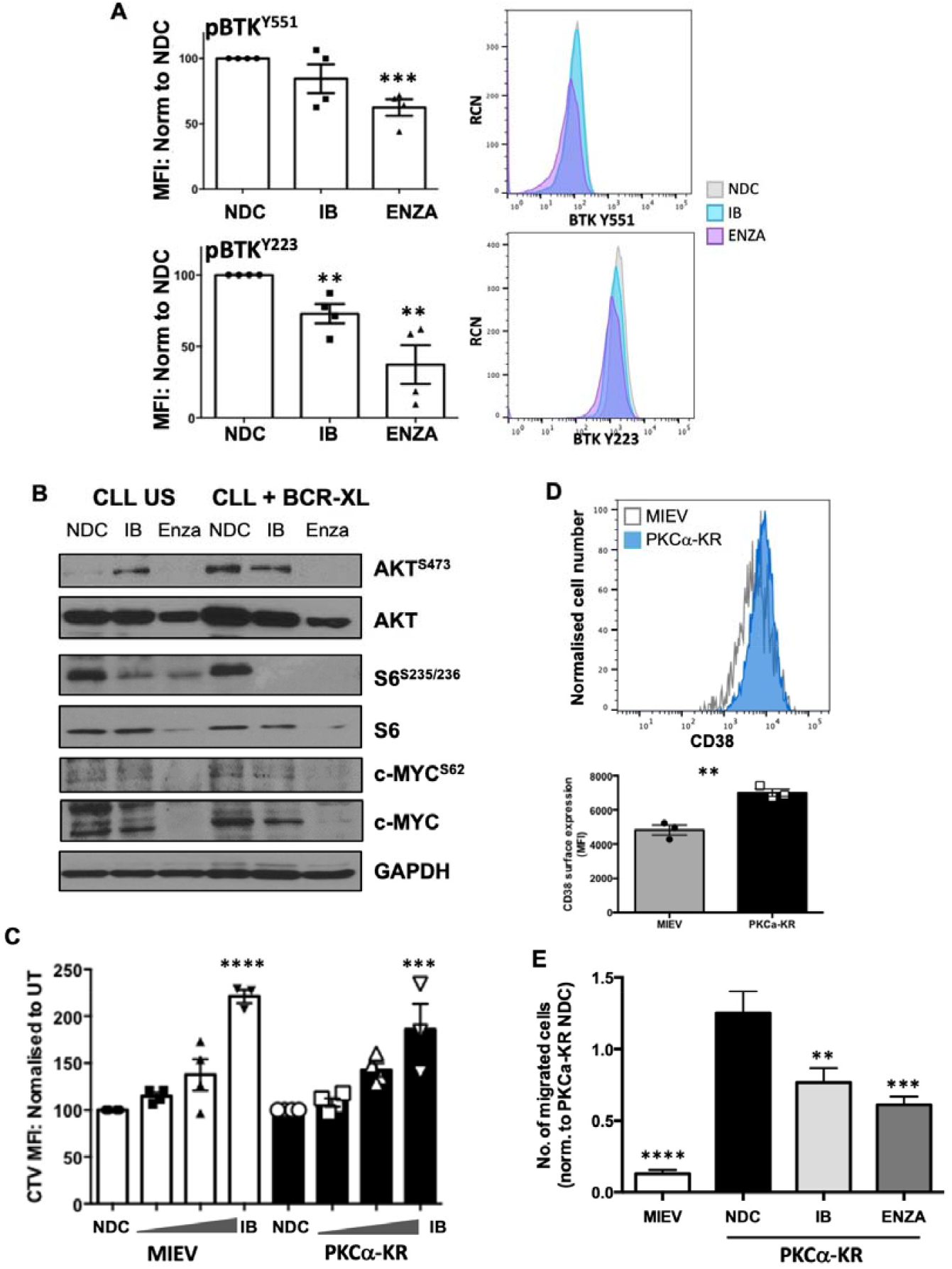
Inhibition of PKCβ upon treatment with enzastaurin downregulates key proteins within the BCR signalosome in primary CLL cells. A & B. Human CLL cells were treated with 1 μM IB or 10 μM Enza in the presence or absence of BCR crosslinking (BCR-XL; F(ab’)_2_ fragment stimulation). A. Phospho-flow was used to analyse the levels of BTK^Y551^ and BTK^Y223^ phosphorylation upon treatment with drugs ± BCR-XL. Left - Graphs show the average MFI of the individual phospho-BTK sites as indicated, normalized to MIEV no drug control (NDC; n=4 individual patient samples). Right - Histogram plots show the effect of drug treatments on BTK^Y551^ and BTK^Y223^ phosphorylation as overlay (NDC – pale grey; IB – turquoise; ENZA – pink); B. Western blots were performed to identify the effect of drug treatment on key proteins within the BCR signalosome in human CLL cells. Representative Western blots are shown. Densitometry of the Western blots for these proteins (n≥3 individual patients) is shown in Supplementary Figure 6; C. Late co-culture MIEV and PKCα-KR cells (2 × 10^6^) were labelled with CTV and cultured for 48 hr in the presence (100 nM – 3 μM) or absence (NDC) of increasing concentrations of IB. Results are expressed as the CTV MFI relative to NDC cells for MIEV and PKCα-KR cultures (n=4 individual experiments); D. A histogram is shown comparing the surface expression of CD38 in MIEV and PKCα-KR cells taken from d33 of co-culture (upper) and the average MFI of CD38 expression is shown relative to MIEV cultures (n=3); E. Migration assessment of MIEV and PKCα-KR cells was performed in the presence and absence of 1 μM IB or 20 μM Enza. The data shown represents an average of 5 independent experiments.

We have previously shown that enza significantly and selectively impacts PKCα-KR cell viability (20 μM) and proliferation (10 μM) over MIEV cells [12]. Testing cell viability with increasing IB concentrations revealed that MIEV cells were more sensitive to IB than PKCα-KR cells up to a concentration of 10 μM IB (Supplementary Figure 7). MIEV and PKCα-KR cells were similarly affected by IB treatment with a significant block in proliferation at 3 μM (Figure 5C). As it has previously been established that inhibition of BCR-mediated signals with IB can reduce CLL cell migration towards CXCL12 [19], and leukocyte migration was the fourth most deregulated pathway in PKCα-KR cells, the migration capacity of MIEV and PKCα-KR CLL-like cells was tested in the presence of IB or Enza treatment. Initial analysis of the adhesion marker CD38, which is regulated by BCR-mediated signaling, showed that surface expression of CD38 was upregulated on PKCα-KR cells compared with MIEV cells (Figure 5D) [20,21]. Our data revealed that PKCα-KR cells have a significantly enhanced migration ability towards SDF1 compared to MIEV cells (Figure 5E), likely due to elevated surface expression of CXCR4 and CD38 (Supplementary Figure 8), and elevated BCR-mediated signaling. However, while migration was significantly inhibited upon treatment with either IB or Enza, these treatments did not alter the surface expression of CD38 (Supplementary Figure 9).

## Discussion

BCR signals are central to CLL pathogenesis, and PKCβ is an important regulator of these signals in healthy and malignant B cells. Although we have known for many years that PKCβII is overexpressed in the malignant cells of CLL [8], the link between PKCβ and leukemogenesis is not yet demonstrated. Here we show that PKCβ expression plays a central role in the development of leukemic cells in our model of CLL (PKCα-KR mouse model). Moreover, we show that the transcription factor SP1 is important for driving a transcription program that promotes leukemogenesis in our model system, and that this program is, at least in part, driven by PKCβ activity. Importantly, this SP1-driven program is also observed in human CLL cells, suggesting a role for PKCβ in the pathogenesis of human disease.

The critical role played by PKCβ in the development of normal B cells has long been established [22]: targeted disruption of the *prkcb* gene in mice results in a severe reduction of marginal zone and B1 populations, while our own work shows that overexpression of PKCβII expands these B cell populations [23]. In contrast, the PKCα knockout mouse model has phenotype in early B lymphocyte development or proliferation and to date, this model has not been associated with the development of hematological malignancies [24]. Indeed, PKCα-deficient mice display an increased risk of developing colorectal cancer which similar to CLL, exhibits a reduction in PKCα expression and an elevation in PKCβII expression in patient samples [25]. In the current study, we demonstrate that both CD5 and CD23 were downregulated in PKCα-KR-transduced cells on a background of *prkcb* KD *in vitro*. This is distinct from the PKCβ-deficient mice, which only exhibited a reduction in CD5^+^ B1 B cells, with surface CD23 expression on B cell populations being unaffected [22]. This indicates that *prkcb* KD targets the leukemic phenotype present in the PKCα-KR mouse model. A relationship between PKCβ and CLL pathogenesis was suggested in studies showing overexpression of this isoform in the malignant cells of this disease, particularly in cells from patients with late stage disease [8]. More recently, PKCβ was found to play a critical role in disease development in the Eμ-TCL1 CLL mouse model, with no leukemia development in the absence of PKCβ expression [10]. Subsequent studies of this model highlighted the essential role played by PKCβII within the stromal compartment in promoting both maintenance and chemosensitivity of malignant B cells [26,27], but did not address whether this PKC isoform was involved in the initiation of malignant cell transformation. Our work provides clarity to this question and shows that PKCβ expression is important for leukemogenesis in our model of CLL, a model we have previously shown is similar to human CLL with respect to overexpression of PKCβII in the leukemic cells [12]. It is likely, however, that the role of PKCβ in this context is one of facilitation rather than transformation because our earlier work has also shown malignant disease does not occur when this PKC isoform is specifically overexpressed in B cells [27]. A similar role for PKCβ overexpression has also been suggested for neoplastic transformation in colon cancer [25,28]. Another BCR pathway protein overexpressed in CLL cells, Lyn, is similar to PKCβ in promoting expansion of the malignant clone through its role within microenvironmental cells, however it differs because Lyn deficiency does not affect the rate of malignant cell transformation in the Eμ-Tcl1 murine model of CLL [29]. Thus, the current study remains the first to directly implicate a signaling protein within the BCR pathway in CLL leukemogenesis.

Our data shows that elevated PKCβ expression in PKCα-KR-transduced cells is driven by elevated Sp1 binding activity at the *prkcb* promoter. This is similar to what is observed in human CLL cells [17] and suggests that the mechanisms controlling PKCβ expression are accurately modelled by our system. Further similarity is demonstrated by our experiments using mtm to target the DNA binding activity of SP1. In addition to *prkcb* downregulation, mtm treatment reduces expression of *blnk, bcl2, vegfa, lef1* and *sp1* in the PKCα-KR-transduced cells, and mtm treatment of CLL cells results in comparable downregulation of the human orthologues of these genes. These data suggest that Sp1, a ubiquitous transcription factor, may play a significant role in promoting a gene transcription profile that induces CLL cell survival, and is aligned with poor prognostic disease [30]. This notion is supported by recent work showing that targeted delivery of miR-29b to CLL cells leads to enhanced cellular reprogramming and decreased viability due to its effect of decreasing SP1 expression [31]. Furthermore, mtm analogs have been shown to be effective against CLL cells, with EC-7072 inducing CLL cell death through targeting of BCR signaling [32], and administration of MTM_OX_32E to the Eμ-Tcl1 mouse model reduces the malignant cell burden [33].

BCR signaling strength in the malignant cells of CLL correlates with poor disease prognosis. This has relevance to our model because it was one of the top deregulated pathways in our microarray comparing gene expression between PKCα-KR-transduced and control cells. We focused on validating this pathway and addressed whether PKCβ activity played a central role in driving this pathway in our mouse model. Biochemical analysis of the activation status of signaling pathways downstream of the BCR indicated that our mouse model possesses constitutively active BCR signaling, as shown by increased expression of key proteins: Lyn, ZAP70, PKCβII, c-Myc and EGR1, together with elevated phosphorylation of BTK^Y223^, AKT^S473^, S6^S235/236^ and c-Myc^S62^. We further noted a downregulation in Lck expression which has previously been described in BCR-activated CLL cells [34]. These validate our microarray data, consolidating and extending our previous findings showing activation of the ERK-MAPK- and AKT/mTOR signaling pathways in PKCα-KR-expressing cells [11,12,35].

The significant upregulation of both BTK^Y551^ and BTK^Y223^ phosphorylation in our mouse model indicates that BTK is constitutively active, leading to an activation of distal BCR-mediated signaling (ie ERK-MAPK pathway), and suggests that such phosphorylation may be aided by elevated Lyn kinase expression which is reported to target phosphorylation of BTK^Y551^ and promote downstream BCR signalling events [36]. A role for PKCβ in regulating this phosphorylation event is shown in experiments using enzastaurin or ibrutinib, which target PKCβ or BTK respectively. We found that both compounds inhibited BTK^Y223^ phosphorylation in PKCα-KR and primary CLL cells, causing reduction in AKT/mTOR and ERK-MAPK/c-Myc mediated pathways. One difference between PKCα-KR and primary CLL cells in this respect is that enzastaurin also inhibited BTK^Y551^ phosphorylation in the latter but not the former. While this result may be related to off-target effects of enzastaurin [37], it may suggest that PKCβII may have an impact of BCR signaling upstream of BTK activation. While the mechanism regulating PKCβII-mediated phosphorylation of BTK is not clear, it is interesting to note that PKCβII has been demonstrated to negatively regulate BTK by S180 phosphorylation to prevent its membrane recruitment [38]. Whether these two events are linked remains to be elucidated in future studies.

While it has been well established that ibrutinib has enhanced the survival of patients with B cell malignancies including CLL, mantle cell lymphoma and Waldenström’s macroglobulinemia (WM) [39-41], clinical trials incorporating enzastaurin have not, as yet, revealed any significant improvement in patient survival in DLBCL [42]. Pre-clinical studies using sotrastaurin, a more broad range PKC isoform inhibitor targeting α, β and θ isoforms, in CLL and DLBCL indicate a strong anti-tumour effect [43,44]. While the reasons for a lack of effect of enzastaurin is not entirely clear, it may be due to the study design as it was used as relapse prevention rather than as disease-active therapy in high risk patients. Given the interest in developing appropriate models to test potential therapeutic compounds, we consider that our CLL-like mouse model provides an excellent model for carrying out pre-clinical testing of novel BCR-targeted agents, that may be translated towards the clinic.

In conclusion, the data reported here demonstrate a role for PKCβ in the development of leukemic B cells in our model, and suggest it may be important for facilitating leukemogenesis in CLL. This has implications for the current understanding of CLL pathogenesis where signaling through the BCR pathway is important. Although many of the kinases in this pathway are overexpressed in CLL cells, PKCβ is the only one currently that facilitates neoplastic transformation of these cells. Moreover, the current study validates our PKCα-KR model for the study of CLL cells where many of the responses we observe using PKCα-KR CLL cells are similar to what is observed in primary CLL cells. Given this parallel, we consider that our CLL-like mouse model will be an excellent tool to decipher the pathobiological behavior of CLL cells.

## Acknowledgements

The authors thank the CLL patients for donating blood samples for this study. This study was funded by a project grant from LLR (Blood Cancer UK – Ref. 13012). Cell sorting facilities were funded by the Kay Kendall Leukaemia Fund (KKL501) and the Howat Foundation. AT was funded by a Bloodwise project grant (13012), AH was supported by a KKLF Clinical training fellowship (KKL838), JH was funded by a Bloodwise (BCUK) project grant (18003), NM was funded by a MRC-DTP PhD studentship.

## Author Contributions

AT & JH designed/performed the majority of experiments, analyzed and interpreted the data, carried out statistical analysis and drafted the manuscript; AH, NM, AFK, IFS, JL, HA performed some experiments and carried out data analysis; AA, HA, BHL carried out the microarray data analysis; KD and JC provided technical assistance with the *in vivo* model and flow cytometry respectively, and analysed the data; NK supervised and designed some the studies; JRS supervised and designed some the studies and wrote the manuscript; AMM obtained funding for the study, designed the research, supervised the studies, analyzed and interpreted the data and wrote the manuscript. All authors reviewed the manuscript.

## References

1. Stevenson FK, Forconi F, Packham G. The meaning and relevance of B-cell receptor structure and function in chronic lymphocytic leukemia. Semin Hematol. 2014;51:158–67.

2. Hamblin TJ, Davis Z, Gardiner A, Oscier DG, Stevenson FK. Unmutated Ig V(H) genes are associated with a more aggressive form of chronic lymphocytic leukemia. Blood. 1999;94(6):1848–54.

3. Burger JA, Chiorazzi N. B cell receptor signaling in chronic lymphocytic leukemia. Trends Immunol. 2013;34:592–601.

4. Slupsky JR. Does B cell receptor signaling in chronic lymphocytic leukemia cells differ from that in other B cell types. Scientifica (Cairo). 2014;2014:208928.

5. Park H, Wahl MI, Afar DE, Turck CW, Rawlings DJ, Tam C, et al. Regulation of Btk function by a major autophosphorylation site within the SH3 domain. Immunity. 1996;4:515–25.

6. Michie AM, Nakagawa R. Elucidating the role of protein kinase C in chronic lymphocytic leukaemia. Hematol Oncol. 2006;24:134–8.

7. Shinohara H, Yasuda T, Aiba Y, Sanjo H, Hamadate M, Watarai H, et al. PKCbeta regulates BCR-mediated IKK activation by facilitating the interaction between TAK1 and CARMA1. J Exp Med. 2005;202:1423–31.

8. Abrams ST, Lakum T, Lin K, Jones GM, Treweeke AT, Farahani M, et al. B-cell receptor signaling in chronic lymphocytic leukemia cells is regulated by overexpressed active protein kinase CbetaII. Blood. 2007;109:1193–201.

9. zum Büschenfelde CM, Wagner M, Lutzny G, Oelsner M, Feuerstacke Y, Decker T, et al. Recruitment of PKC-betaII to lipid rafts mediates apoptosis-resistance in chronic lymphocytic leukemia expressing ZAP-70. Leukemia. 2010;24:141–52.

10. Holler C, Piñón JD, Denk U, Heyder C, Hofbauer S, Greil R, et al. PKCbeta is essential for the development of chronic lymphocytic leukemia in the TCL1 transgenic mouse model: validation of PKCbeta as a therapeutic target in chronic lymphocytic leukemia. Blood. 2009;113:2791–4.

11. Nakagawa R, Soh JW, Michie AM. Subversion of PKCa signaling in hematopoietic progenitor cells results in the generation of a B-CLL-like population in vivo. Cancer Research. 2006;66:527–34.

12. Nakagawa R, Vukovic M, Tarafdar A, Cosimo E, Dunn K, McCaig AM, et al. Generation of a poor prognostic chronic lymphocytic leukemia-like disease model: PKCα subversion induces an upregulation of PKCβII expression in B lymphocytes. Haematologica. 2015;100:499–510.

13. McCaig AM, Cosimo E, Leach MT, Michie AM. Dasatinib inhibits B cell receptor signalling in chronic lymphocytic leukaemia but novel combination approaches are required to overcome additional pro-survival microenvironmental signals. Br J Haematol. 2011;153:199–211.

14. Cosimo E, McCaig AM, Carter-Brzezinski LJ, Wheadon H, Leach MT, Le Ster K, et al. Inhibition of NF-κB-mediated signaling by the cyclin-dependent kinase inhibitor CR8 overcomes prosurvival stimuli to induce apoptosis in chronic lymphocytic leukemia cells. Clin Cancer Res. 2013;19:2393–405.

15. Harris WJ, Huang X, Lynch JT, Spencer GJ, Hitchin JR, Li Y, et al. The histone demethylase KDM1A sustains the oncogenic potential of MLL-AF9 leukemia stem cells. Cancer Cell. 2012;17:473–87.

16. Tarafdar A, Dobbin E, Corrigan P, Freeburn R, Wheadon H. Canonical Wnt signaling promotes early hematopoietic progenitor formation and erythroid specification during embryonic stem cell differentiation. PLoS One. 2013;8:e81030.

17. Al-Sanabra O, Duckworth AD, Glenn MA, Brown BR, Angelillo P, Lee K, et al. Transcriptional mechanism of vascular endothelial growth factor-induced expression of protein kinase CβII in chronic lymphocytic leukaemia cells. Sci Rep. 2017;7:43228.

18. Ray R, Snyder RC, Thomas S, Koller CA, Miller DM. Mithramycin blocks protein binding and function of the SV40 early promoter. J Clin Invest. 1989;83:2003–7.

19. de Rooij MFM, Kuil A, Geest CR, Eldering E, Chang BY, Buggy JJ, et al. The clinically active BTK inhibitor PCI-32765 targets B-cell receptor- and chemokine-controlled adhesion and migration in chronic lymphocytic leukemia. Blood. 2012;119:2590–4.

20. Zupo S, Isnardi L, Megna M. Massara R, Malavasi F, Dono M, et al. CD38 expression distinguishes two groups of B-cell chronic lymphocytic leukemias with difference responses to anti-IgM antibodies and propensity to apoptosis. Blood. 1996;88:1365.

21. Lanham S, Hamblin T, Oscier D, Ibbotson R, Stevenson F, Packham G. Differential signaling via surface IgM is associated with VH gene mutational status and CD38 expression in chronic lymphocytic leukemia. Blood. 2003;101:1087.

22. Leitges M, Schmedt C, Guinamard R, Davoust J, Schaal S, Stabel S, et al. Immunodeficiency in protein kinase cbeta-deficient mice. Science. 1996;273:788–91.

23. Azar AA, Michie AM, Tarafdar A, Malik N, Menon GK, Till KJ, et al. A novel transgenic mouse strain expressing PKCβII demonstrates expansion of B1 and marginal zone B cell populations. Sci Rep. 2020;10:13156.

24. Pfeifhofer C, Gruber T, Letschka T, Thuille N, Lutz-Nicoladoni C, Hermann-Kleiter N, et al. Defective Ig2a/2b class switching in PKC alpha-/-mice. J Immunol. 2006;176:6004.

25. Gökmen-Polar Y, Murray NR, Velasco MA, Gatalica Z, Fields AP. Elevated protein kinase C betaII is an early promotive event in colon carcinogenesis. Cancer Res. 2001;61:1375–81.

26. Lutzny G, Kocher T, Schmidt-Supprian M, Rudelius M, Klein-Hitpass L, Finch AJ, et al. Protein kinase c-β-dependent activation of NF-κB in stromal cells is indispensable for the survival of chronic lymphocytic leukemia B cells in vivo. Cancer Cell. 2013;23:77–92.

27. Park E, Chen J, Moore A, Mangolini M, Santoro A, Boyd JR, et al. Stromal cell protein kinase C-β inhibition enhances chemosensitivity in B cell malignancies and overcomes drug resistance. Sci Transl Med. 2020;12:eaax9340.

28. Murray NR, Davidson LA, Chapkin RS, Clay Gustafson W, Schattenberg DG, Fields AP. Overexpression of protein kinase C betaII induces colonic hyperproliferation and increased sensitivity to colon carcinogenesis. J Cell Biol. 1999;145:699–711.

29. Nguyen PH, Fedorchenko O, Rosen N, Koch M, Barthel R, Winarski T, et al. LYN kinase in the tumor microenvironment is essential for the progression of chronic lymphocytic leukemia. Cancer Cell. 2016;30:610–22.

30. Wu W, Zhu H, Fu Y, Shen W, Miao K, Hong M, et al. High LEF1 expression predicts adverse prognosis in chronic lymphocytic leukemia and may be targeted by ethacrynic acid. Oncotarget. 2016;7:21631–43.

31. Chiang CL, Goswami S, Frissora FW, Xie Z, Yan PS, Bundschuh R, et al. ROR1-targeted delivery of miR-29b induces cell cycle arrest and therapeutic benefit in vivo in a CLL mouse model. Blood. 2019;134:432–44.

32. Lorenzo-Herrero S, Sordo-Bahamonde C, Bretones G, Payer ÁR, González-Rodríguez AP, González-García E, et al. The Mithralog EC-7072 Induces Chronic Lymphocytic Leukemia Cell Death by Targeting Tonic B-Cell Receptor Signaling. Front Immunol. 2019;10:2455.

33. Rivas JR, Liu Y, Alhakeem SS, Eckenrode JM, Marti F, Collard JP, et al. Interleukin-10 suppression enhances T-cell antitumor immunity and responses to checkpoint blockade in chronic lymphocytic leukemia. Leukemia. 2021; Online ahead of print.

34. Schleiss C, Carapito R, Fornecker LM, Muller L, Paul N, Tahar O, et al. Temporal multiomic modeling reveals a B-cell receptor proliferative program in chronic lymphocytic leukemia. Leukemia. 2021;35:1463–74.

35. Cosimo E, Tarafdar A, Moles MW, Holroyd AK, Malik N, Catherwood MA, et al. AKT/mTORC2 inhibition activates FOXO1 function in CLL cells reducing B cell receptor-mediated survival. Clin Cancer Res. 2019;25(5):1574–87.

36. Wahl MI, Fluckiger AC, Kato RM, Park H, Witte ON, Rawlings DJ. Phosphorylation of two regulatory tyrosine residues in the activation of Bruton’s tyrosine kinase via alternative receptors. Proc Natl Acad Sci U S A. 1997;94:11526–33.

37. Graff JR, McNulty AM, Hanna KR, Konicek BW, Lynch RL, Bailey SN, et al. The protein kinase Cbeta-selective inhibitor Enzastaurin (LY317615.HCl), suppresses signalling through the AKT pathway, induces apoptosis, and suppresses growth of human colon cancer and glioblastoma xenografts. Cancer Res. 2005;65:7462–9.

38. Kang SW, Wahl MI, Chu J, Kitaura J, Kawakami Y, Kato RM, et al. PKCbeta modulates antigen receptor signaling via regulation of Btk membrane localization. EMBO J. 2001;20:5692–702.

39. Coutré SE, Furman RR, Flinn IW, Burger JA, Blum K, Sharman J, et al. Extended Treatment with Single-Agent Ibrutinib at the 420 mg Dose Leads to Durable Responses in Chronic Lymphocytic Leukemia/Small Lymphocytic Lymphoma. Clin Cancer Res. 2017;23:1149–55.

40. Treon SP, Tripsas CK, Meid K, Warren D, Varma G, Green R, et al. Ibrutinib in previously treated Waldenstrom’s macroglobulinemia. N Engl J Med. 2015;372(15):1430–40.

41. Wang ML, Rule S, Martin P, Goy A, Auer R, Kahl BS, et al. Targeting BTK with ibrutinib in relapsed or refractory mantle-cell lymphoma. N Engl J Med. 2013;369(6):507–16.

42. Crump M, Leppä S, Fayad L, Lee JJ, Rocco AD, Ogura M, et al. Randomized, Double-Blind, Phase III Trial of Enzastaurin Versus Placebo in Patients Achieving Remission After First-Line Therapy for High-Risk Diffuse Large B-Cell Lymphoma. J Clin Oncol. 2016;34:2484–92.

43. El-Gamal D, Williams K, LaFollette TD, Cannon M, Blachly JS, Zhong Y, et al. PKC-β as a therapeutic target in CLL: PKC inhibitor AEB071 demonstrates preclinical activity in CLL. Blood. 2014;124:1481–91.

44. Naylor TL, Tang H, Ratsch BA, Enns A, Loo A, Chen L, et al. Protein kinase C inhibitor sotrastaurin selectively inhibits the growth of CD79 mutant diffuse large B-cell lymphomas. Cancer Res. 2011;71:2643–53.

