## Supplemental Data for "PKCβ facilitates leukemogenesis in chronic lymphocytic leukaemia by promoting constitutive BCR-mediated signaling"

**Supplementary Table 1. CLL patient clinical characteristics and their use in the paper.**

^a^ If previously undergone treatment, it was not within three months of sample collection.

^b^ ZAP-70 analysis was conducted by immunohistochemistry in the regional haematology laboratory. ND – not determined.

| **CLL ID** | **Treatment^a^** | **Sex** | **Binet Stage** | **ZAP-70 status^b^** | **Cytogenetics** |
| --- | --- | --- | --- | --- | --- |
| 85 | Yes | F | A | ND | del(11q) |
| 86 | No | F | A | pos | del(11q) |
| 91 | Yes | M | C | pos | del(11q) |
| 93 | Yes | M | C | pos | del(17p) |
| 102 | Yes | F | C | pos | no del(11q) or del(17p) |
| 113 | Yes | F | C | high | del(17p) |
| 144 | No | M | B | low | del(17p) |
| 148 | Yes | M | B | low | del(11q) |
| 150 | No | M | A | high | no del(11q) or del(17p) |
| 151 | No | M | B | ND | del(11q) |

**Supplementary Table 2. Flow Cytometry Antibodies.**

| Name | Conjugation | Clone | Company |
| --- | --- | --- | --- |
| CD45 | Per-CP | 30-F11 | BDBiosciences |
| CD19 | APC-Cy7 | 1D3 | BDBiosciences |
| B220 | PE | RA3-6132 | BDBiosciences |
| CD5 | APC | 53-7.3 | BDBiosciences |
| CD5 | BV510 | 53-7.3 | BioLegend |
| CD23 | PE-Cy7 | B3B4 | BioLegend |
| CXCR4 | BV510 | 2B11 | BDBiosciences |
| CD49d | AF647 | R1-2 | BioLegend |
| CD38 | APC-FIRE | 90 | BioLegend |
| CD38 | PE | 90 | BDBiosciences |
| BTK^Y223^ | AF647 | N35-86 | BDBiosciences |
| BTK^Y551^ | AF647 | 24a/BTK | BDBiosciences |
| Annexin V | APC | N/A | BDBiosciences |

**Supplementary Table 3. PCR primer sequences.**

All sequences are shown 5’ - 3”.

| **Gene** | **Forward sequence** | **Reverse sequence** |
| --- | --- | --- |
| *Btk* | cac cag aaa gac aga ttc cg | cca tag cat tct tgg ctg tc |
| *Egr1* | gag atg atg ctg ctg agc aa | gtc gtt tgg ctg gga taa ct |
| *Tbp* | gta ccc ttc acc aat gac tc | cag cca aga ttc acg gta ga |

**Supplementary Table 4. ChIP primer sequences for the Sp1 binding sites on the mouse *prkcb* promotor region.**

| **SP1 binding sites** | **Forward sequence** | **Reverse sequence** |
| --- | --- | --- |
| *Prkcb* prom 1 | gcg ttt ggt cat tgc tgg at | aca cac aca tac acg tac acc g |
| *Prkcb* prom 2 | cgg tgt acg tgt atg tgt gtg t | ccc tca ttt gca tga aac cc |
| *Prkcb* prom 3 | tgt ctg tgt gtg tct ctg ct | tgg tcc agc tgt gct tgg ca |

Supplementary Table 5: List of antibodies used for Western blotting.

List of antibodies (Ab) and their dilutions used in TBS-T with milk or BSA. All antibodies were purchased from Cell Signalling Technologies (CST), Santa-Cruz, BD Biosciences or Abcam.

| Name | Clone | Dilution | 2^nd^ary Ab | Company |
| --- | --- | --- | --- | --- |
| PKCα | #2056 | 1:1000 | Rabbit | CST |
| pAKT^S473^ | D9E | 1:1000 | Rabbit | CST |
| AKT (pan) | C67E7 | 1:1000 | Rabbit | CST |
| pS6^S235/S236^ | D57.2.2E | 1:1000 | Rabbit | CST |
| S6 | 54D2 | 1:1000 | Mouse | CST |
| PKCβII | C-18 | 1:1000 | Rabbit | Santa-Cruz |
| SP1 | ab227383 | 1:500 | Rabbit | Abcam |
| LYN | #2732 | 1:1000 | Mouse | CST |
| LCK | 28/Lck | 1:1000 | Mouse | BDBiosciences |
| EGR1 | 44D5 | 1:1000 | Rabbit | CST |
| SYK | #2712 | 1:1000 | Rabbit | CST |
| c-MYC^S62^ | EPR17924 | 1:1000 | Rabbit | Abcam |
| c-Myc | #9402 | 1:1000 | Rabbit | CST |
| GAPDH | D16H11 | 1:1000 | Rabbit | CST |
| α-mouse IgG, HRP Ab | #7076 | 1:10000 | Horse | CST |
| α-rabbit IgG, HRP Ab | #7074 | 1:10000 | Goat | CST |

**Supplementary Table 6**: CLL cells (1 x 10^7^ cells) from 4 patient samples were purified from whole blood and then incubated ± 200 nM mithramycin. Following treatment, RNA was harvested from the cells, assessed for quality, and subjected to gene expression analysis using a Whole human genome (4x44K) microarray kit (Agilent Technologies). The effect of mithramycin treatment is recorded as fold change in gene expression together with the adjusted P value for the 4 patient samples analyzed. The microarray data are available at the GEO repository (GSE210348).

| Gene name | Fold change | Adjusted P value |
| --- | --- | --- |
| PRKCB | -3.70 | p=0.00022 |
| BCL2 | -1.92 | p=0.00031 |
| LEF1 | -1.47 | p=0.00032 |
| BLNK | -1.09 | p=0.0020 |
| VEGFA | -0.561 | p=0.011 |
| SP1 | -0.529 | P=0.041 |

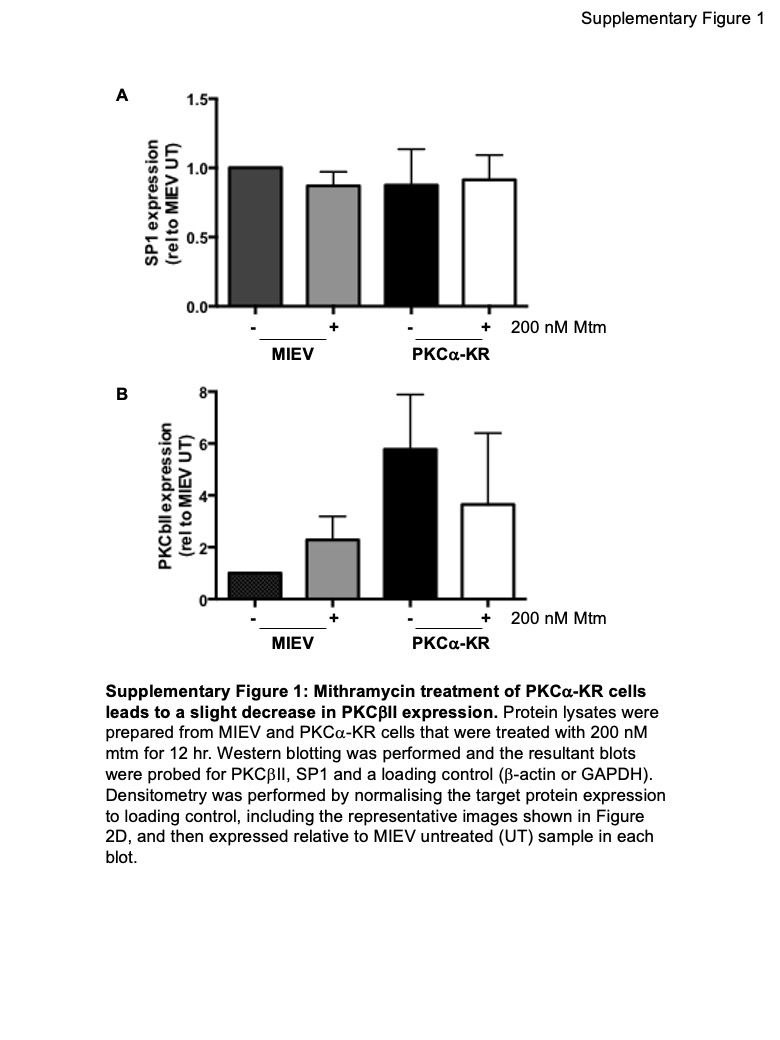

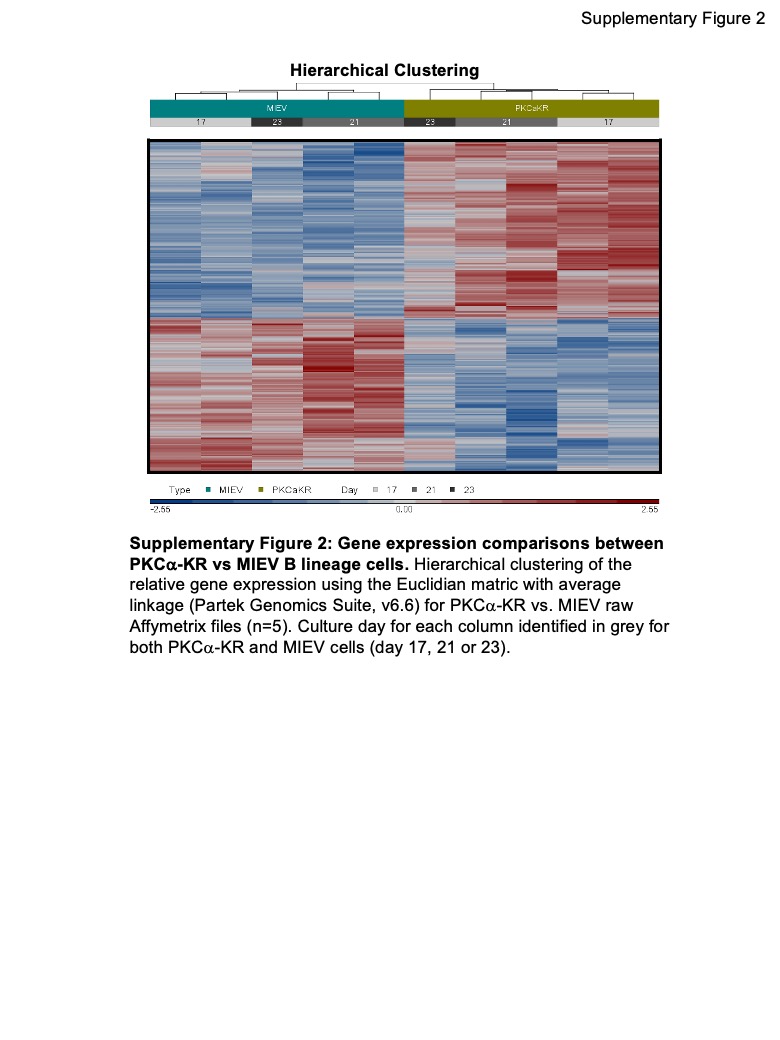

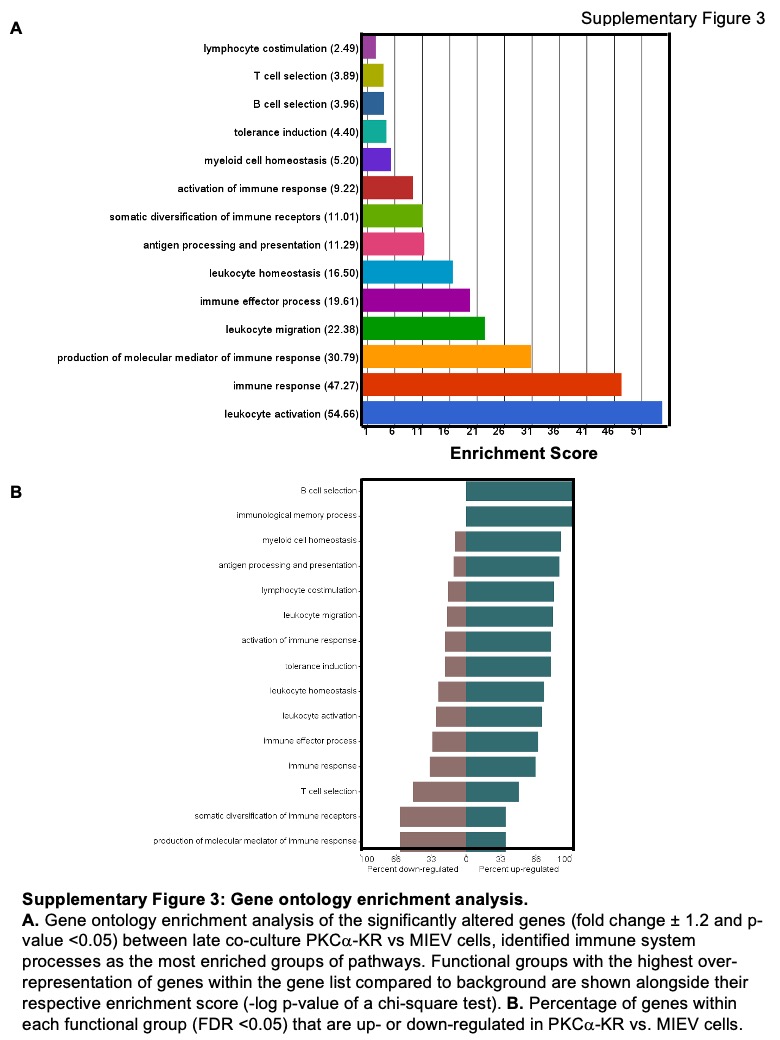

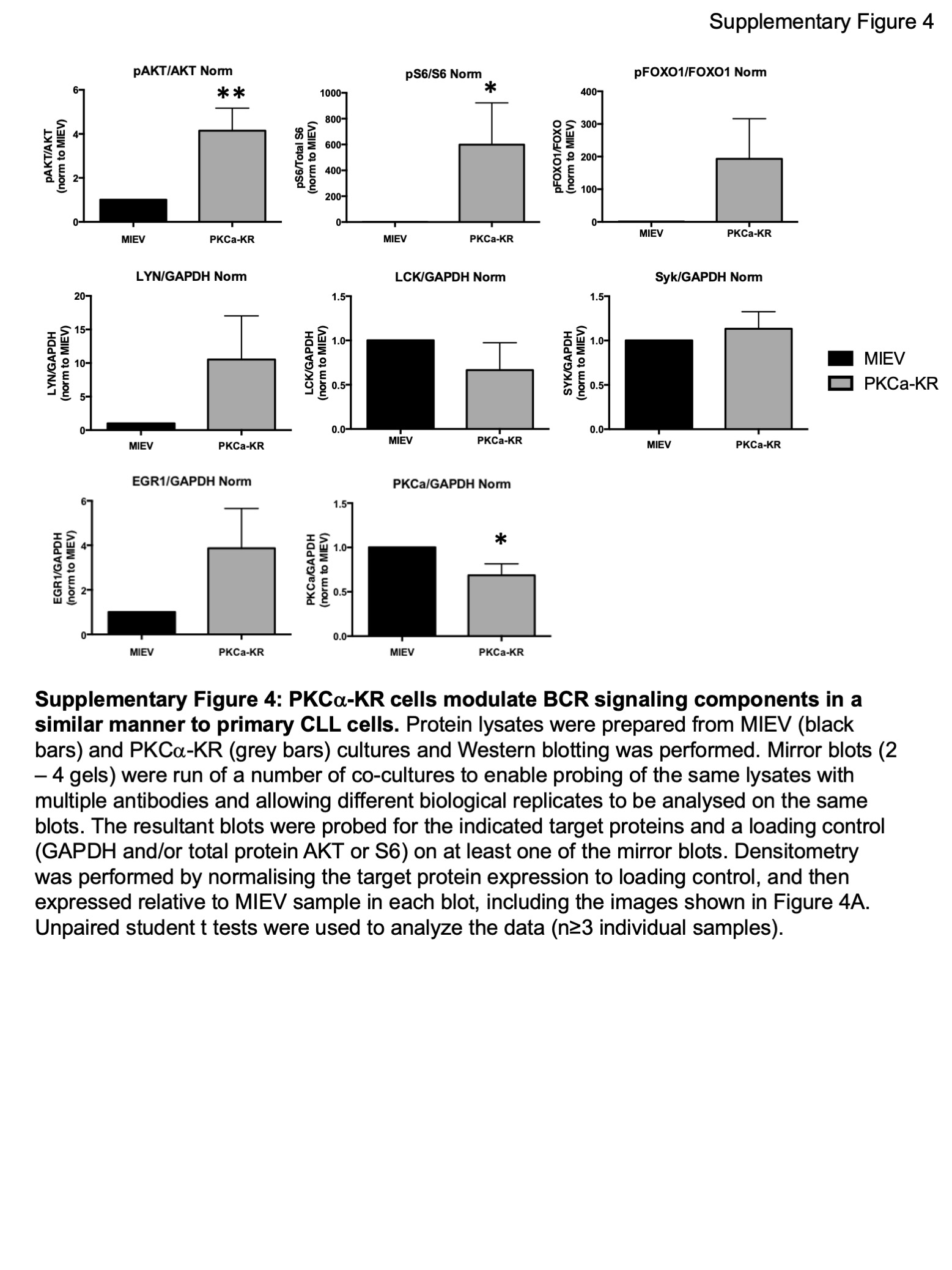

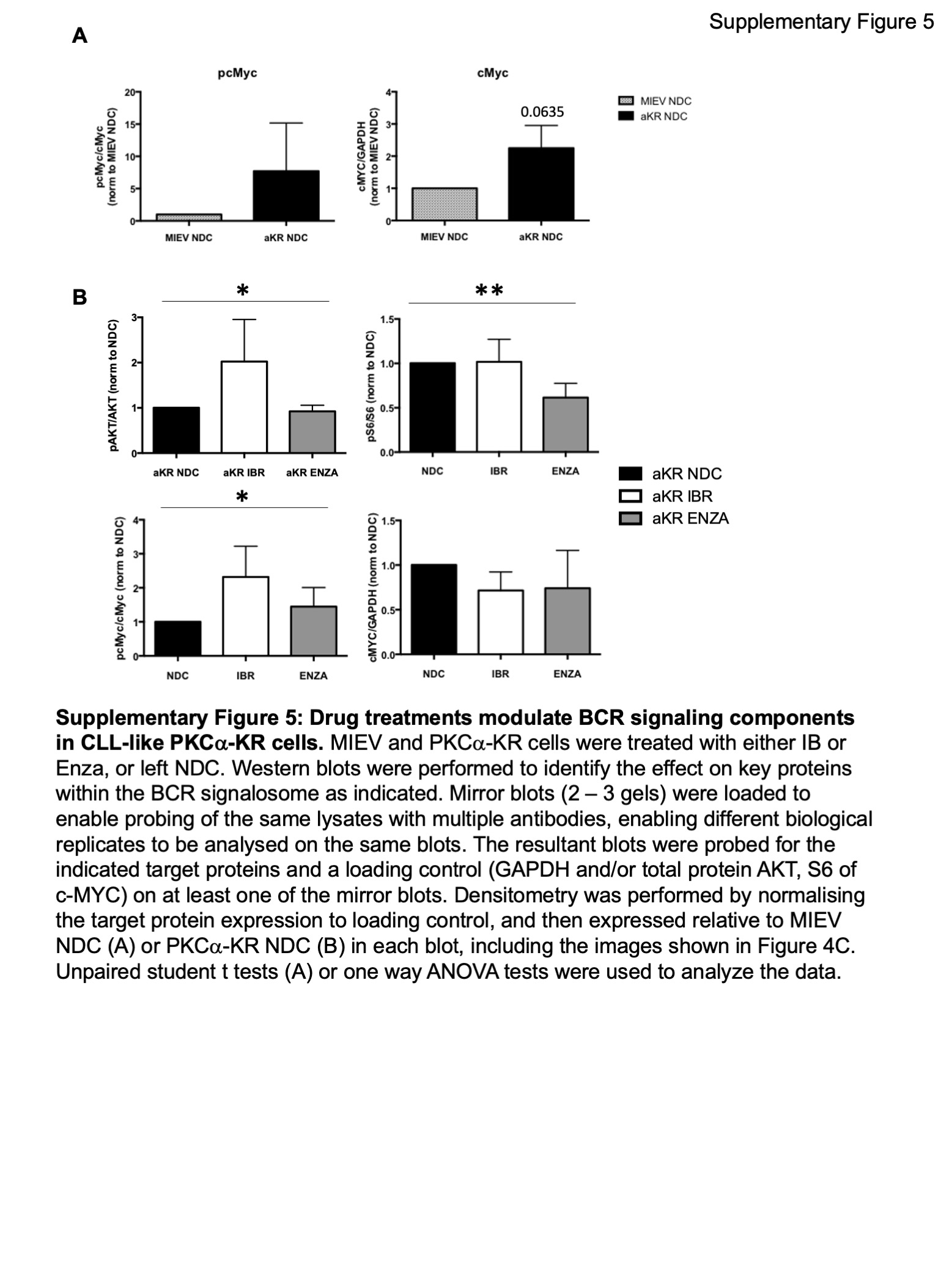

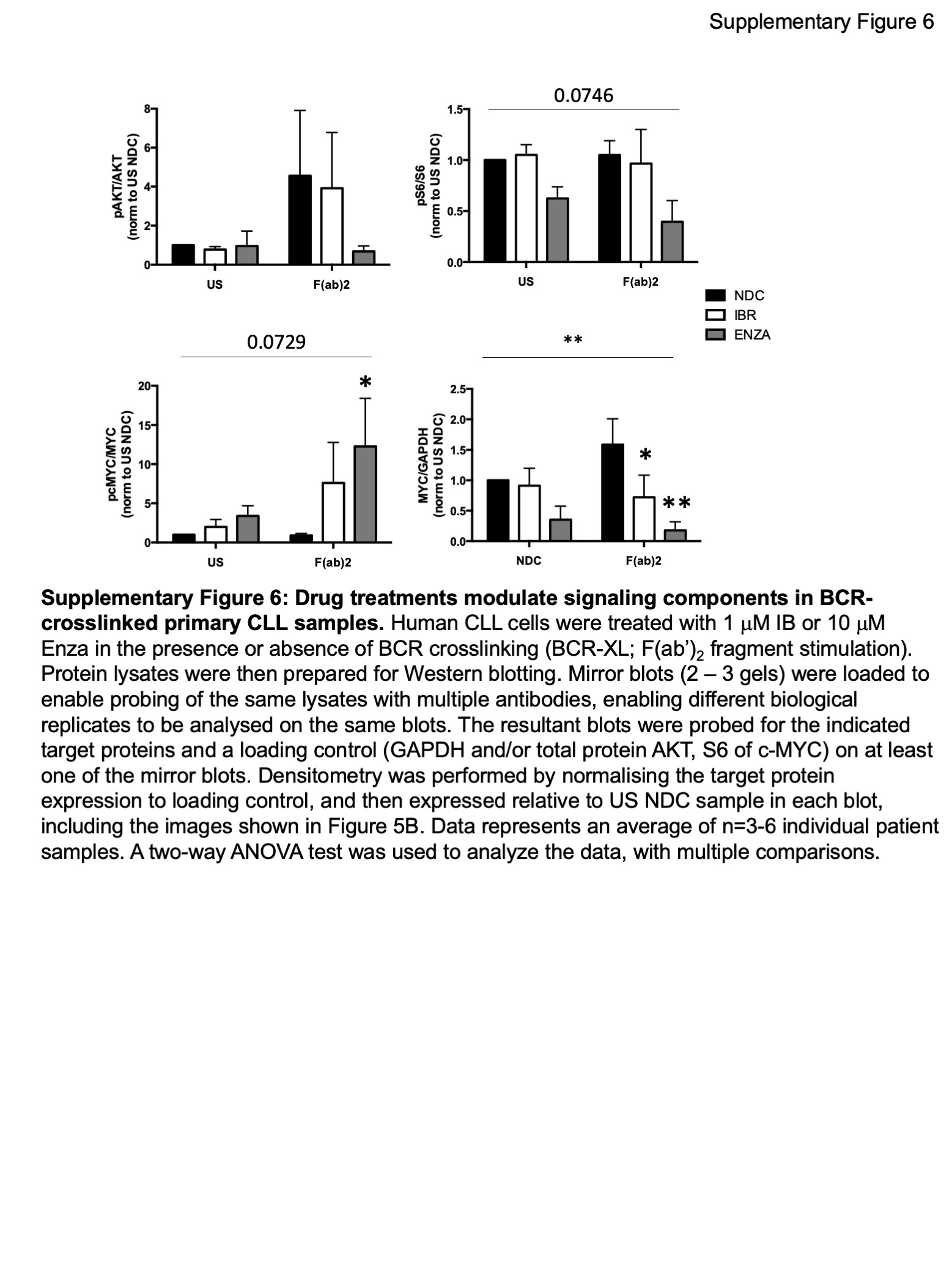

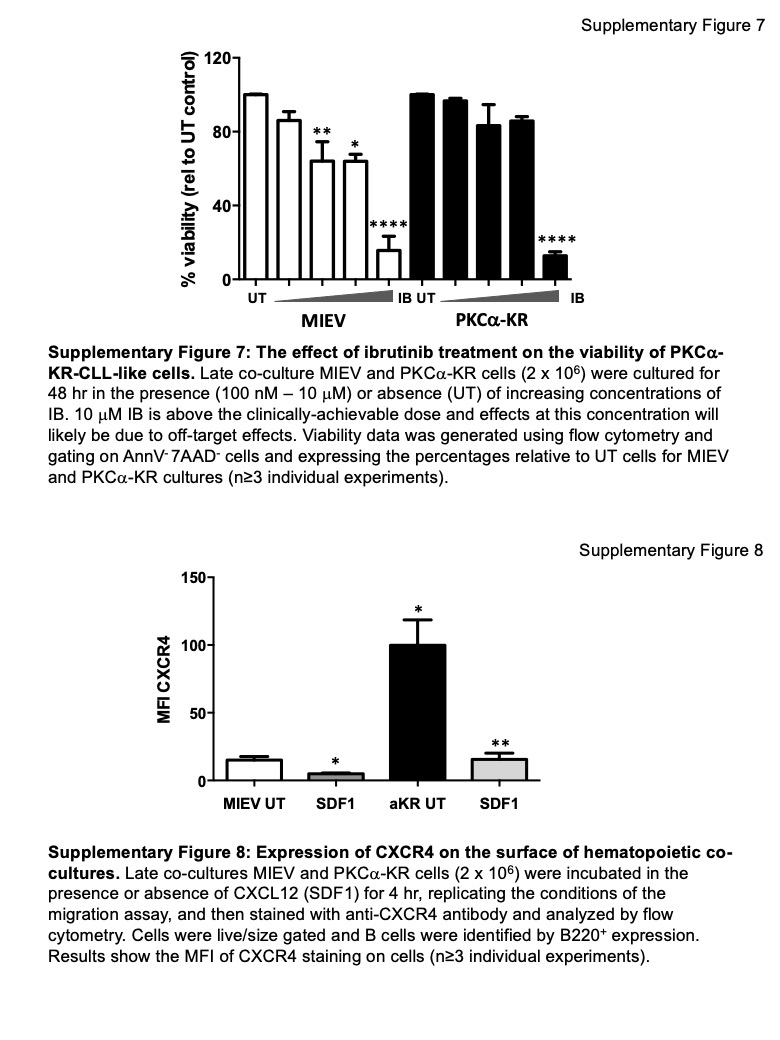

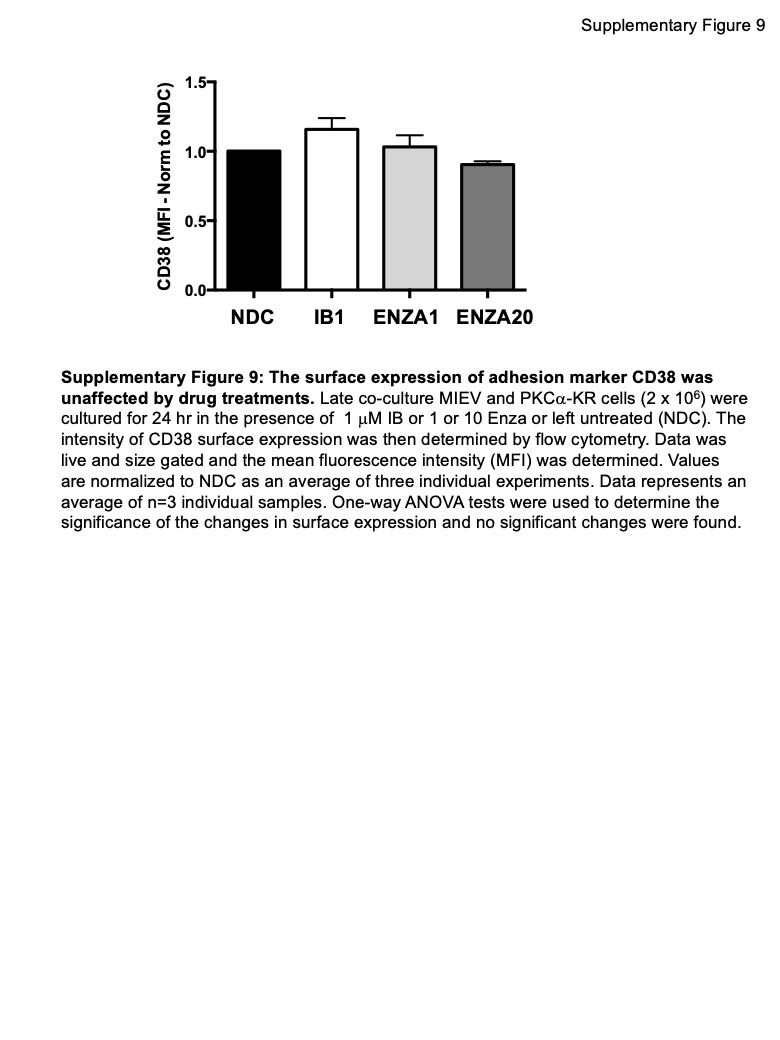

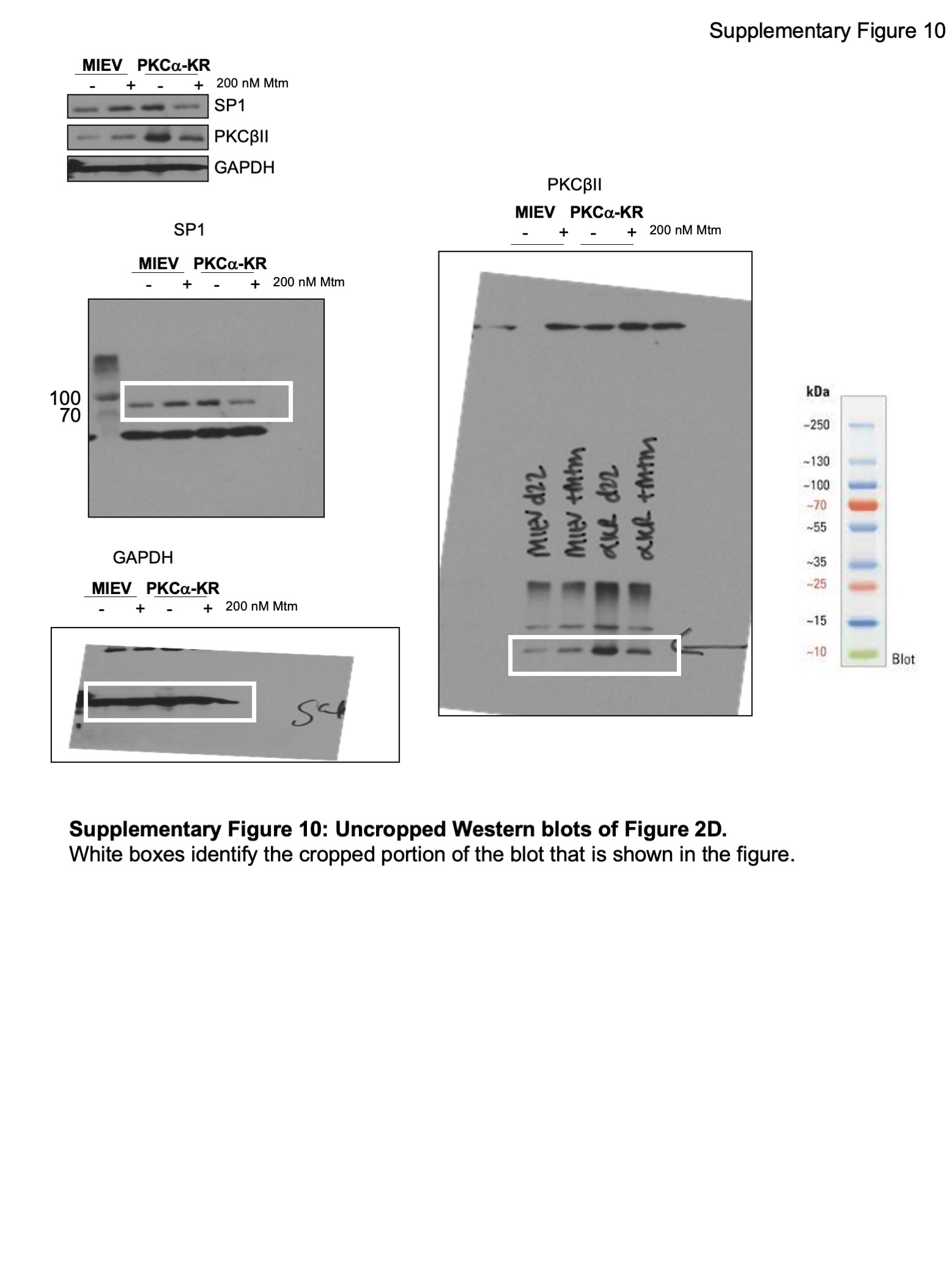

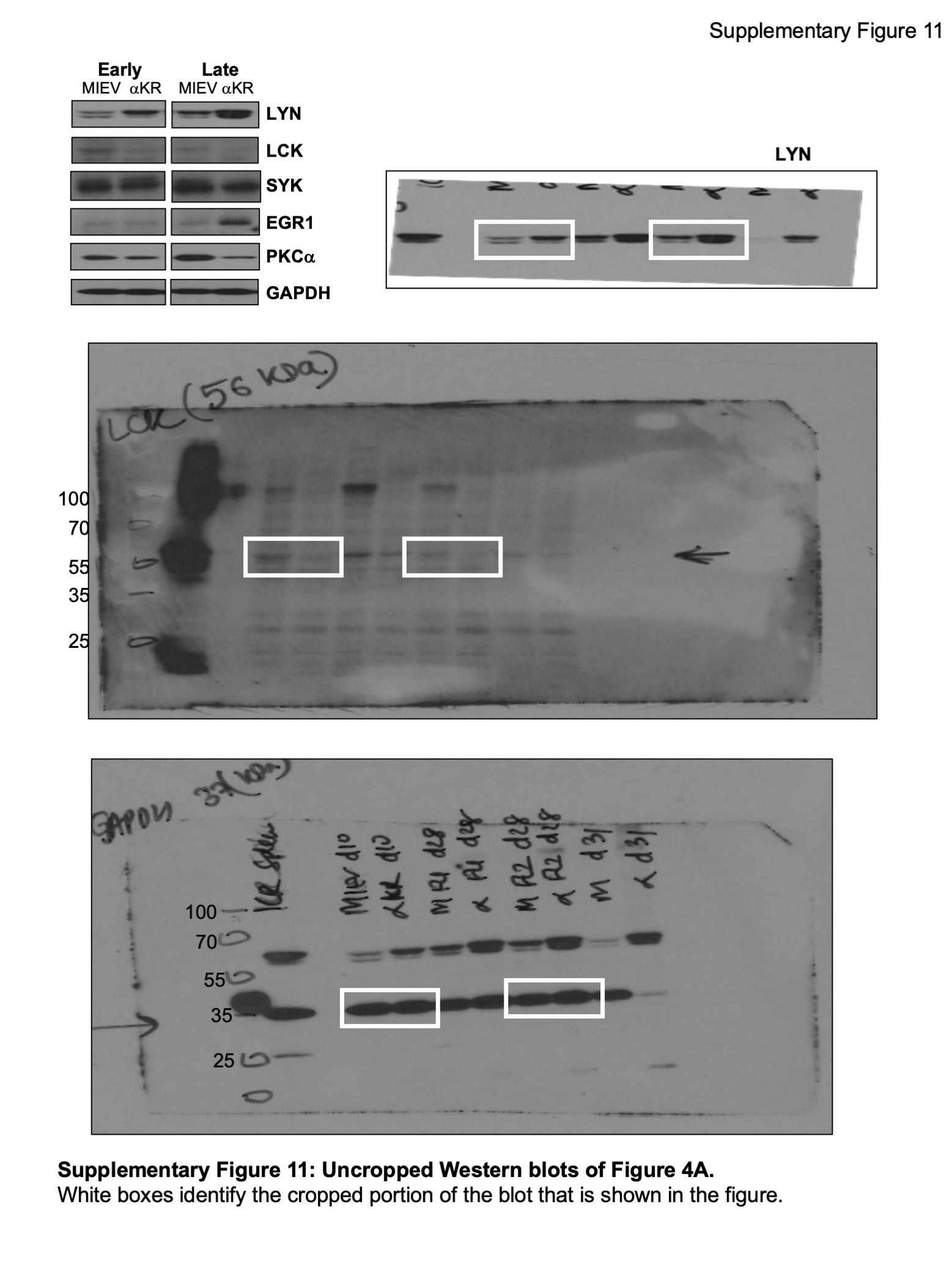

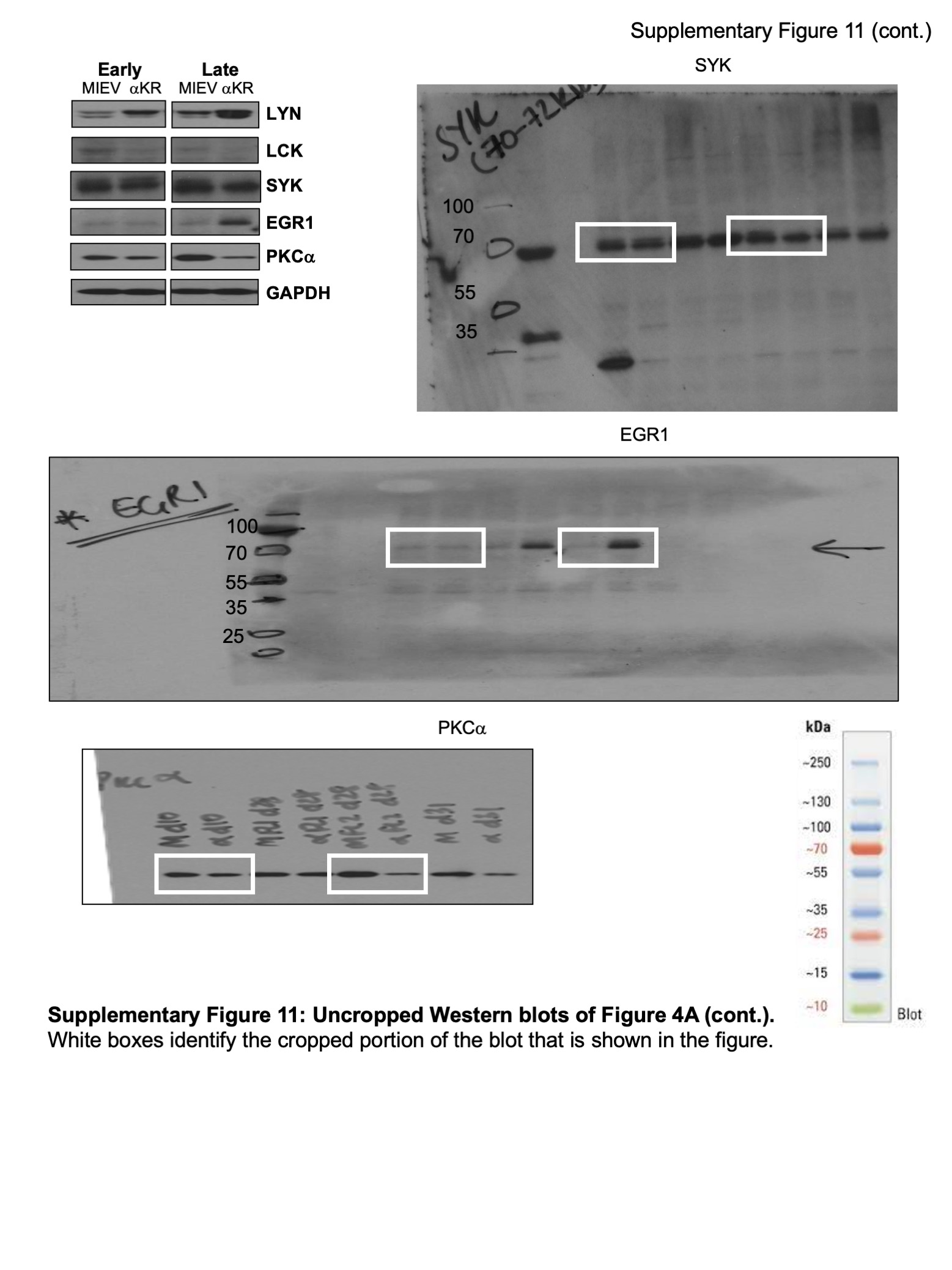

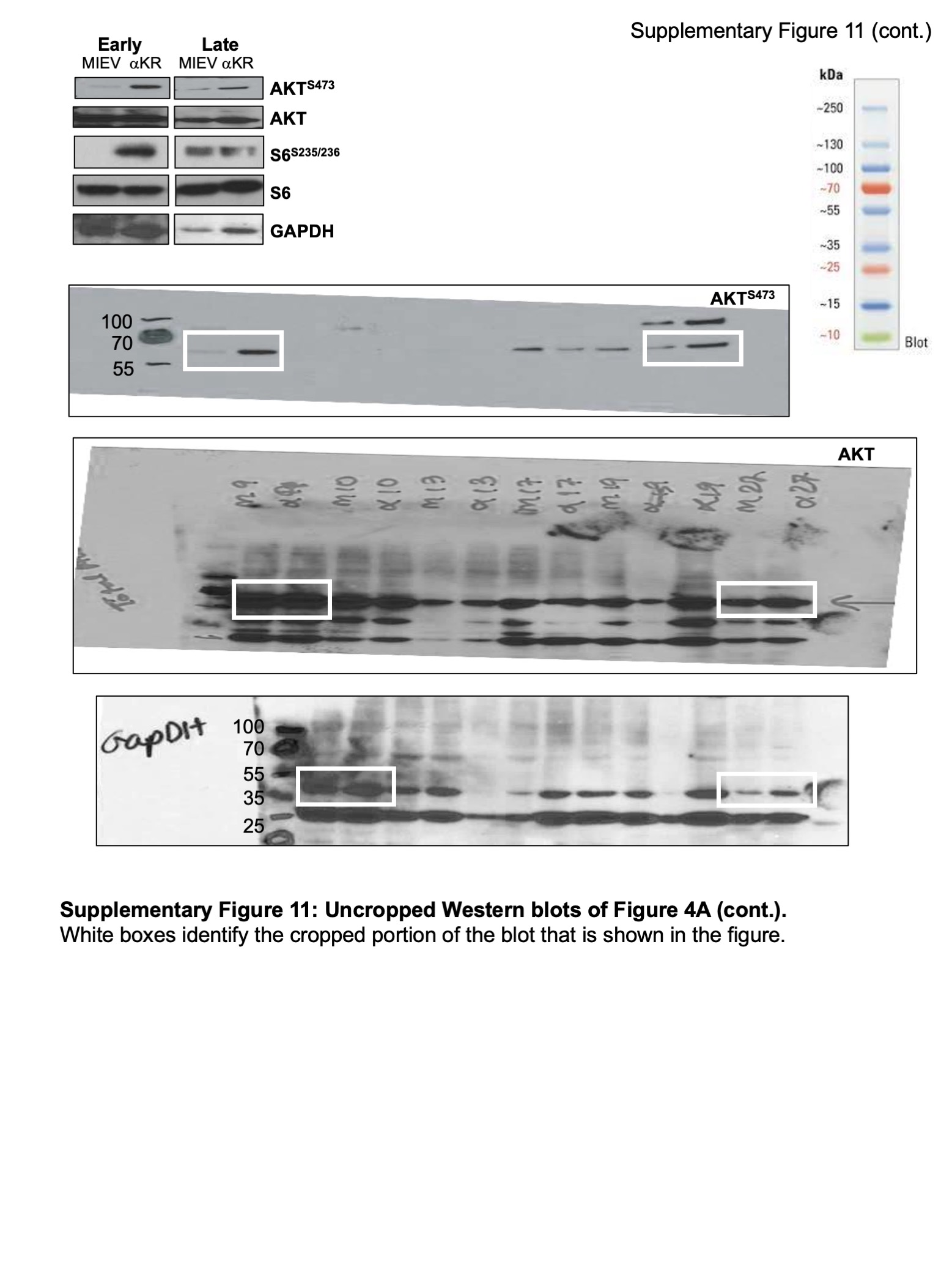

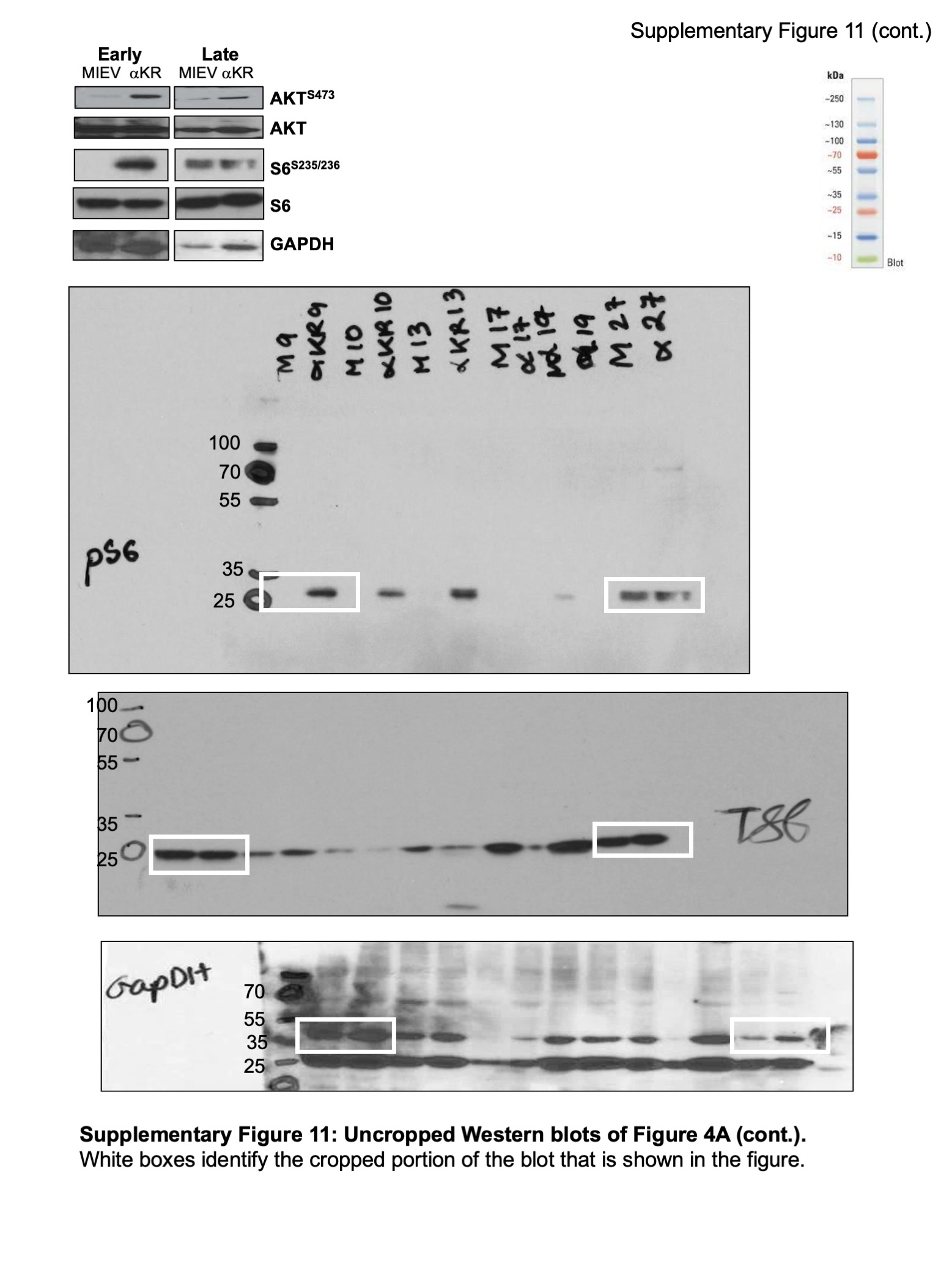

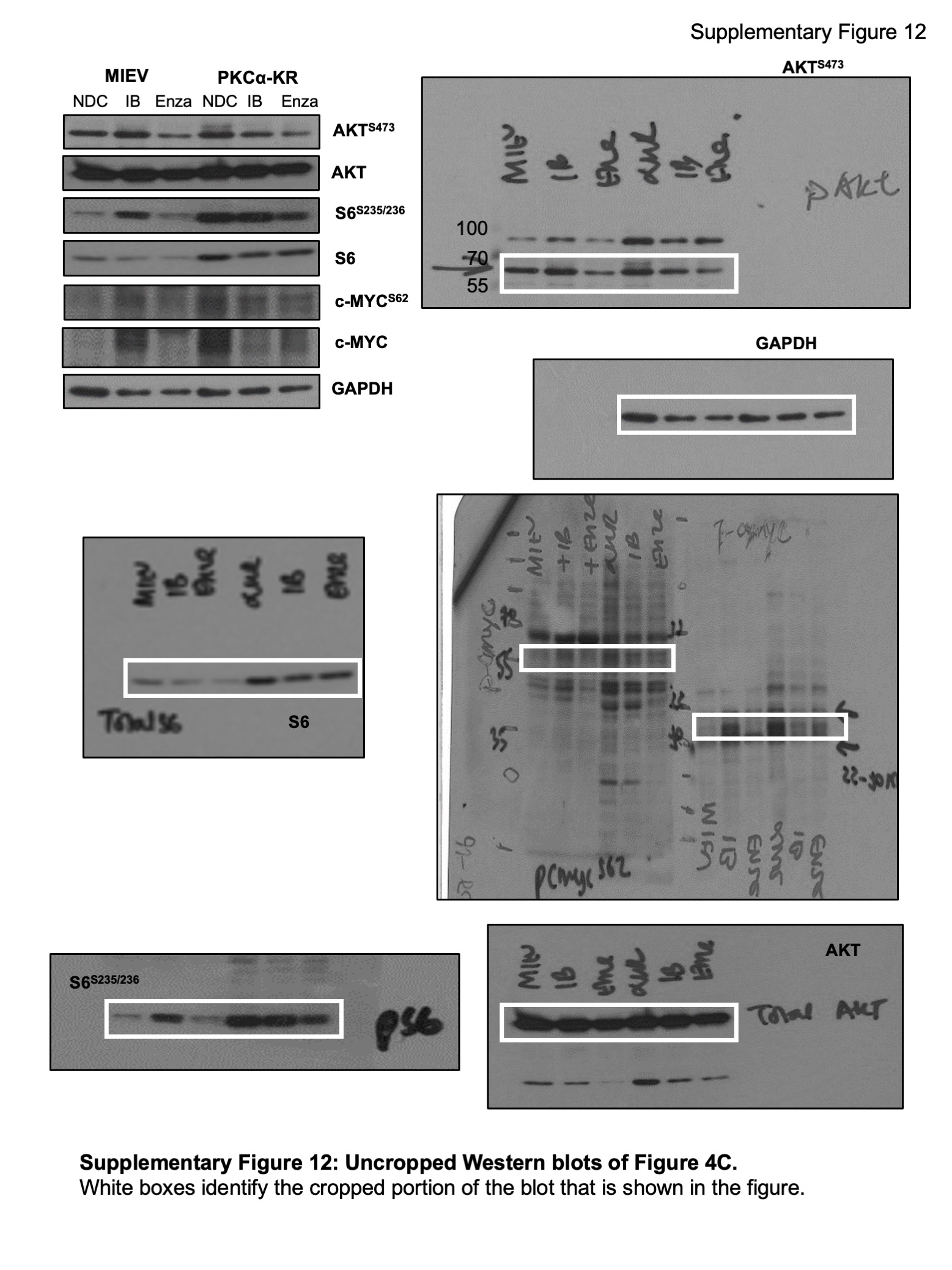

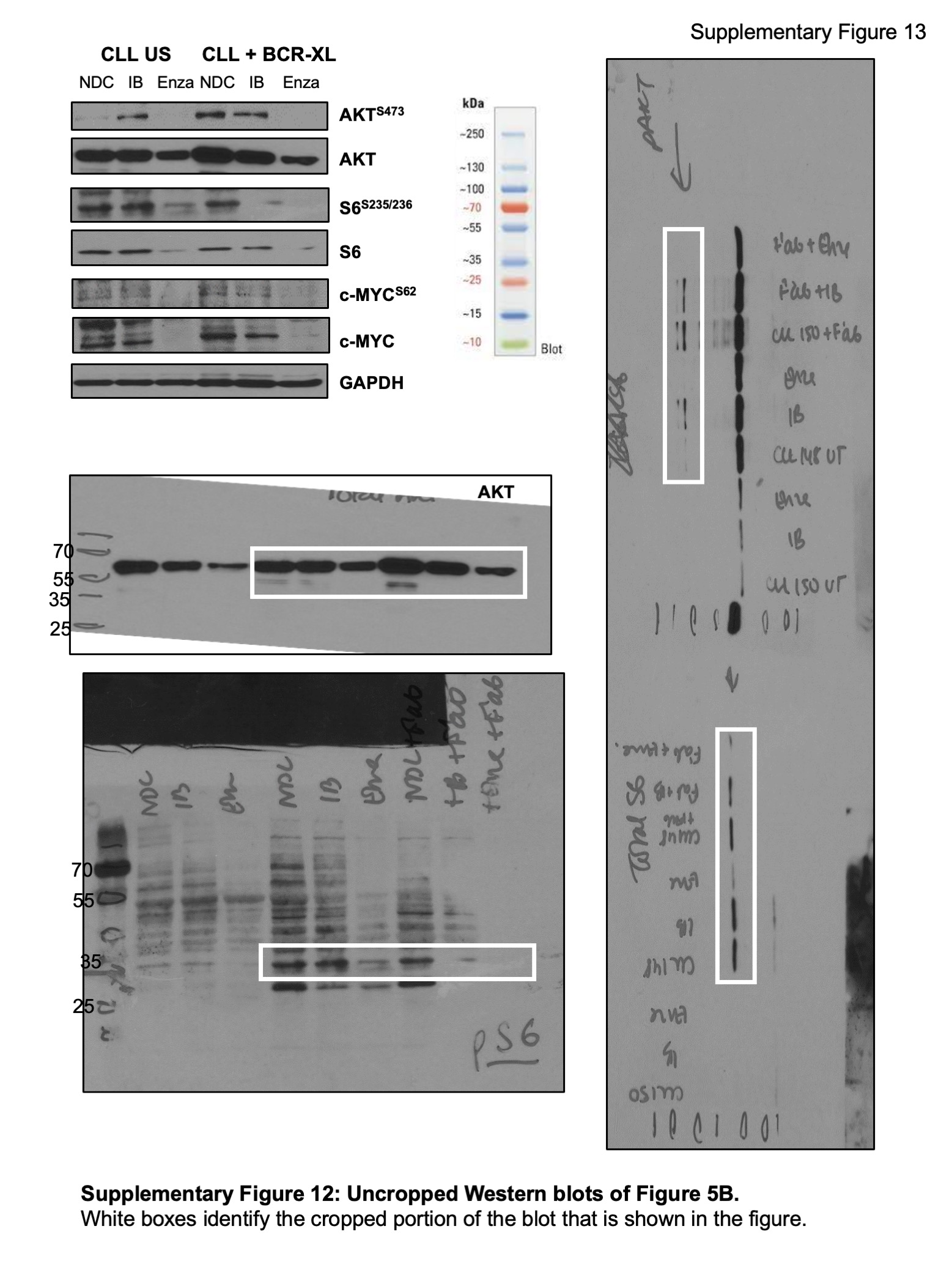

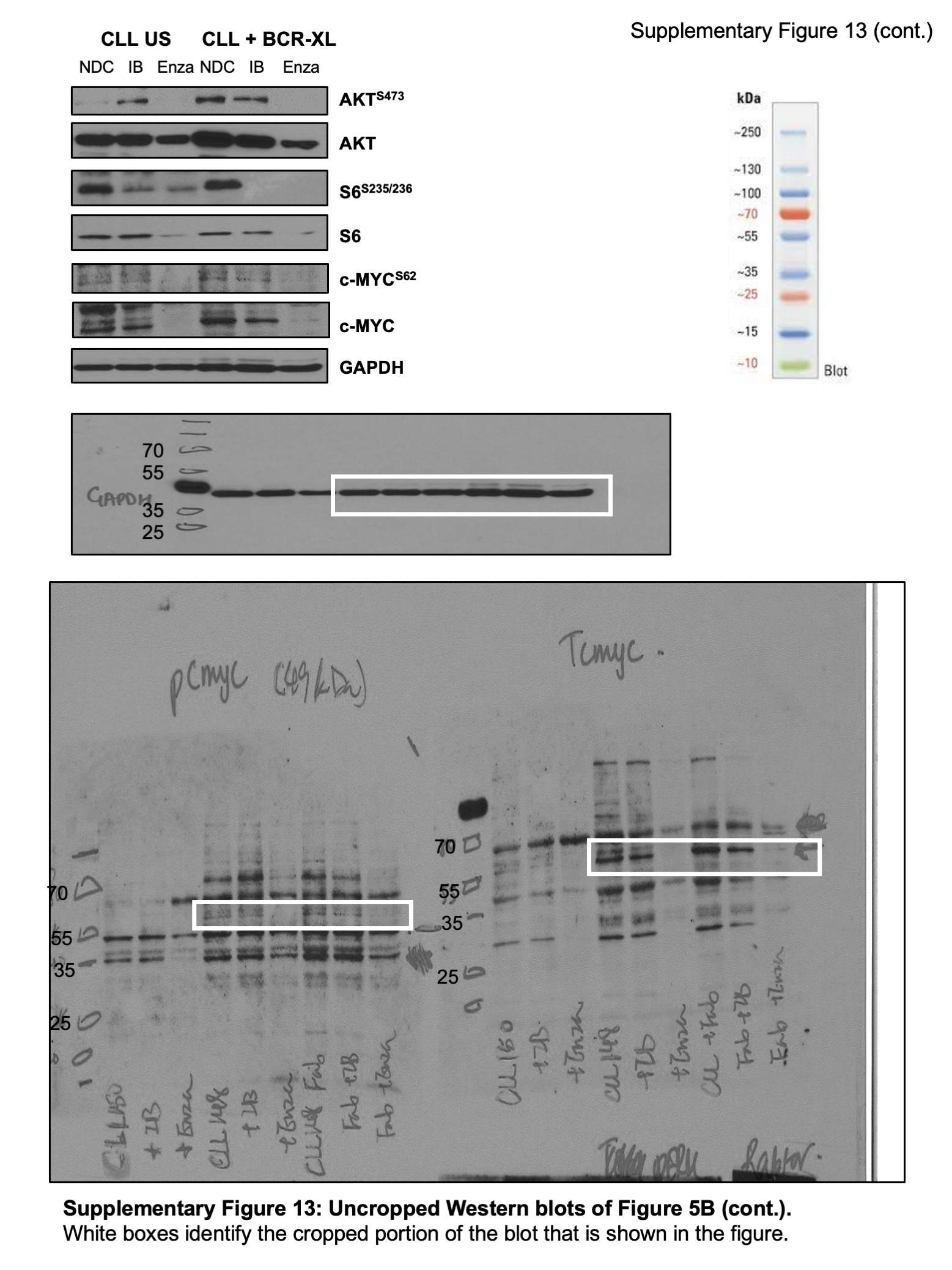
